# Transcription factors organize into functional groups on the linear genome and in 3D chromatin

**DOI:** 10.1101/2022.04.06.487423

**Authors:** Rakesh Netha Vadnala, Sridhar Hannenhalli, Leelavati Narlikar, Rahul Siddharthan

## Abstract

Transcription factors (TFs) and their binding sites have evolved to interact cooperatively or competitively with each other. Here we examine in detail, across multiple cell lines, such cooperation or competition among TFs both in sequential and spatial proximity (using chromatin conformation capture assays) on one hand, and based on both in vivo binding as well as TF binding motifs on the other. We ascertain significantly co-occurring (“attractive”) or avoiding (“repulsive”) TF pairs using robust randomized models that retain the essential characteristics of the experimental data. Across human cell lines TFs organize into two groups, with intra-group attraction and inter-group repulsion. This is true for both sequential and spatial proximity, as well as for both in vivo binding and motifs. Attractive TF pairs exhibit significantly more physical interactions suggesting an underlying mechanism. The two TF groups differ significantly in their genomic and network properties, as well in their function—while one group regulates housekeeping function, the other potentially regulates lineage-specific functions, that are disrupted in cancer. We also show that weaker binding sites tend to occur in spatially interacting regions of the genome. Our results suggest a complex pattern of spatial cooperativity of TFs that has evolved along with the genome to support housekeeping and lineage-specific functions.

## 1 Introduction

DNA-binding proteins play a key role in nuclear and cellular processes. In particular, RNA polymerase transcribes genes to RNA molecules; transcription factors (TFs) help regulate transcription by binding to specific sites in DNA; histones organize DNA into a three-dimensional chromatin structure; and the cohesin complex and the CTCF factor further control this 3D organization. Gene expression is controlled by the complicated combinatorial interplay of accessibility of DNA, remodelling of chromatin, binding of proteins such as TFs to DNA, recruitment of RNA polymerase, and the spatial interaction of all these factors on a three-dimensional stage.

All these aspects can be probed today by high-throughput techniques: Hi-C and variants can detect interacting regions in chromatin [1, 2, 3, 4, 5]; ChIP-seq and related techniques can assay binding sites for proteins in vivo genome-wide [6]; and open chromatin can be ascertained by techniques such as DNase-seq and ATAC-seq [7, 8].

Transcriptional regulation has traditionally been studied by considering DNA as a linear sequence containing genes, promoters, enhancers and TF-binding sequence motifs [9, 10, 11, 12, 13]. Only recently have attempts been made to integrate multiple sources of data to form a three-dimensional understanding of transcriptional regulation. Malin et al.[14] showed that spatially clustered enhancers containing low affinity homotypic TF motif sites exhibit greater in vivo TF occupancy than the motif sites not appearing in clusters, highlighting the role of spatial enhancer cluster in potentially assisting binding at weaker TF binding sites. Ma et al. [15] studied co-localization of homotypic and heterotypic motif sites in Hi-C contact regions, suggesting the existence of a spatial TF interaction network. Others have considered 3D chromatin in the context of protein-protein interaction (PPI) networks and pathways[16, 17, 18].

Here, using chromatin interaction (ChIA-PET) and ChIP-seq data in multiple cell lines, as well as motif information, we explore the interplay between TFs in three-dimensional chromatin. Specifically, we examine the co-binding of pairs of TFs in spatially proximal regions and assess whether such co-binding occurs significantly more or less often than one would expect by chance. This is accomplished with a careful randomization technique that respects essential properties of the interaction network.

While our method differs from Ma et al.[15], our predictions largely agree with theirs in GM12878 cells for the TFs that they consider. We consider many more TFs, across multiple cell lines and multiple data sets. We additionally include histone marks associated with various chromatin states in this analysis. We perform and compare with similar analyses for sequentially-adjacent binding and for co-occurrence of motifs in spatially adjacent sequence.

In the cell lines examined here, we find that almost all pairs of TFs tend to co-occur either significantly more often (which we call “attraction”), or significantly less often (which we call “repulsion”) than by chance. For a given cell line, we find that the pattern of attraction or repulsion agrees with what is seen in sequentially-adjacent regions. Interestingly, a similar pattern emerges if one uses only motif information, i.e. no experimental binding data. These two groups appear functionally different, in their location of binding relative to transcriptional start sites (TSS), in their protein-protein-interaction (PPI) network properties, and in the functional nature of their target genes.

We provide a tool to perform similar analyses on chromatin interaction data (such as Hi-C or ChIAPET) and ChIP-seq (or motif) datasets, ChromTogether available at https://github.com/rakeshnetha14/ChromTogether. In the future we hope to use such information to improve the performance of motif-finding and in-silico TFBS prediction.

## Materials and Methods

### Approach

High-throughput chromatin conformation capture-based experiments (such as Hi-C, ChIA-pet, capture Hi-C) report an interaction score between multiple pairs of regions in the genome. We use ChIA-pet data and consider all reported interactions. After merging regions that occur within 2kbp of each other and filtering out pairs of interactions within 2kbp of each other, we obtain a set of long-range interaction pairs. We represent this data as an undirected graph, with a node denoting a region and an edge denoting a long-range interaction. We remove regions with an unrealistically high degree (>20) or length (>30kbp) as these are likely the result of the merging step or from the experiment reporting a cell population average. We call this the “interaction network” arising from that experiment.

We next look at TF-binding information from ChIP-seq data for the same cell-type. We construct a corresponding “binding network” which contains all region nodes from the interaction network as well as additional TF nodes. An edge in this network connects a TF node to a region node, if there is evidence of the TF binding to that region in a ChIP-seq experiment. The interaction network and binding network are illustrated in Figure 1A and 1B.

**Figure 1:**
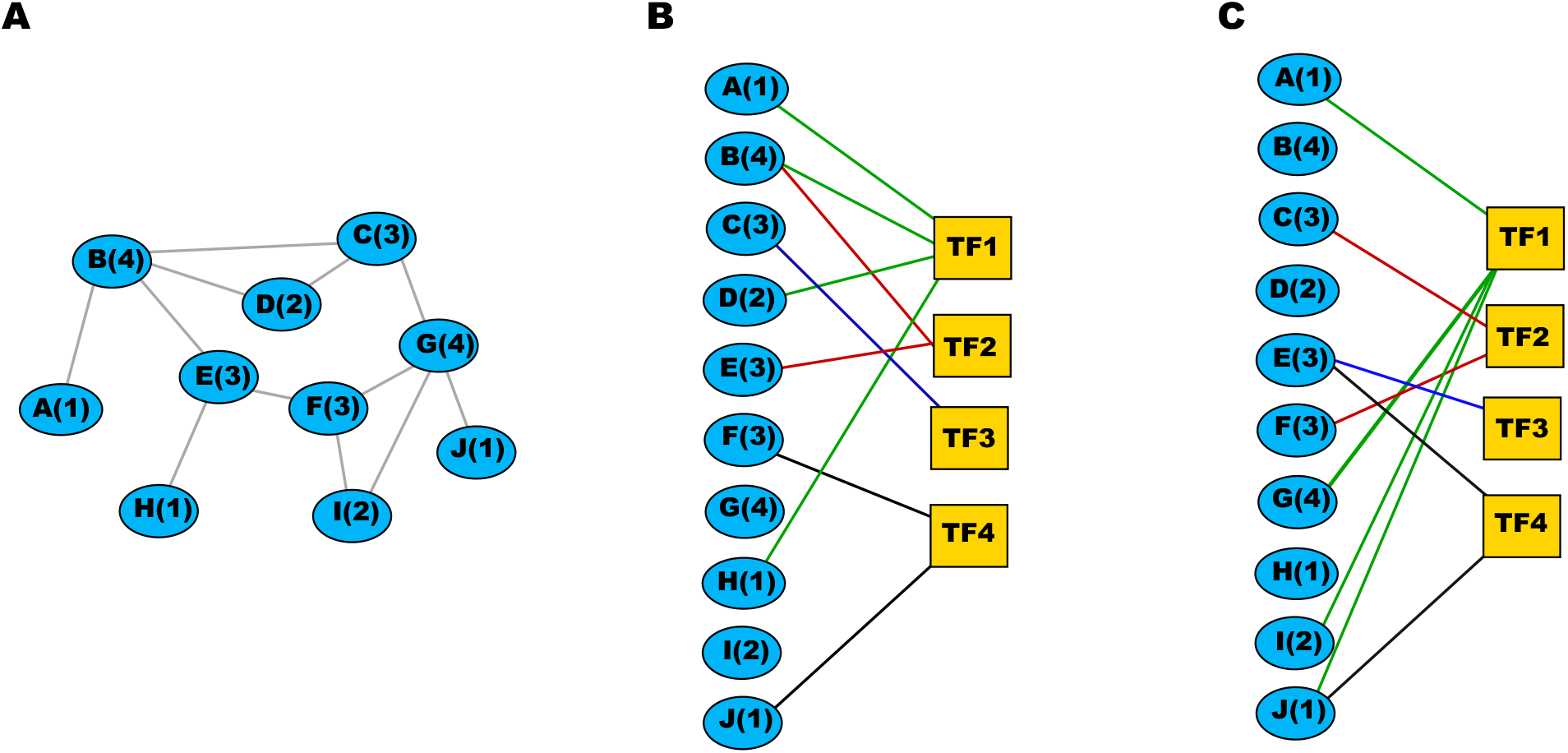
(a) The interaction network, where each node is a contiguous genomic region and links indicate contacts between regions as determined by chromatin interaction data. The degree of each region in the region-region interaction network is shown in brackets. (b) The binding network, a bipartite network where blue nodes are regions as in (a), yellow squares are TFs, and links indicate binding of a TF to a region. Links from TF1 are shown in green, from TF2 in red, for clarity. (b) A possible randomization of links where each TF-region link is randomly reassigned from the TF to another (possibly the same) region such that the region-region interaction degree of the bound region, in brackets, is approximately preserved.

The goal is to assess TF pairs that co-occur via the interaction network. For this purpose, all instances of TFs *i* and *j* binding to adjacent nodes in the interaction network are considered as co-occurrences of the two TFs. We estimate the significance of these co-occurrences by simulating random networks whose construction is described below. Defining co-occurrence that is significantly more frequent than random as “attraction” and co-occurrence that is significantly less frequent than random as “repulsion”, we ask whether there is any biological significance to attracting or repelling pairs; whether such pairs are conserved across cell types and across species; whether such attracting (repelling) pairs also attract (repel) within short (500–2000bp) contiguous regulatory sequence (eg, promoters or enhancers); and whether, if one looks purely at sequence motifs rather than ChIP-seq binding events, similar patterns are seen.

Our randomized networks are constructed to preserve essential properties of the original data. We view the binding network as a bipartite network, with nodes representing regions in the interaction network linked to nodes representing TFs in the ChIP-seq data. The region-region interaction degree of each region is known and the node corresponding to each region is labelled, in the binding network, with its region-region interaction degree. We generate random networks by reassigning these TF-region links with the constraint that each TF-region link is assigned from the same TF to a region with a similar region-region degree. Specifically: we reassign a TF link to a region *x* of degree ≤ 5 to a random region *y* with exactly the same region-region degree as region *x*; a region *x* with degree between 5 and 10 to a random region y with degree within ± 2 of region *x*’s degree; and a region *x* with degree >10 to a random region of degree >10. This ensures that in the randomized bipartite network, each TF node has the same out-degree as originally, and that the interaction-network degree distribution for TF target regions is approximately conserved. This is illustrated in figure 1 (C). The supplementary material figure S1 shows this conservation for each TF-region in the original network throughout the 1000 randomization steps.

We construct 1000 such randomized networks. From these, we calculate two *p*-values for each pair of TFs: *p_ij_* is the *p*-value for TFs *i* and *j* co-occurring equally or more often than in the real data under the null that they co-occur randomly, calculated as the fraction of random networks where this happens; and similarly 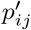 is the *p*-value for *i* and *j* co-occurring equally or *less* often than in the real data. To correct for multiple pairwise comparisons, we use the full list of *p*-values to calculate *q*-values *q* and *q′*, which indicate false discovery rates, following the Benjamini-Hochberg procedure [19]. Small values of *q*, and *q′*, respectively indicate TF pairs co-occurring significantly more often, and less often, than chance. Finally, we show the significant *q* and *q′* values (less than 0.05) on a heatmap, in green and red respectively. In order to show both attraction and repulsion on the same heatmap, we plot *q* in green and 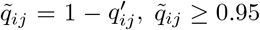 in red (with the brightest reds being the largest, i.e. most significant values). As a control, we selected one of the random network as real network and compared it with all other random networks and results in no significant attraction or repulsion TF pairs.

The chromatin interaction datasets and uniformly processed ChIP-seq peaks generated by ENCODE project used in the study are given in the supplementary tables S1 and S2.

For motif analysis, position weight matrices (PWMs) were taken from the JASPAR database[20]. When multiple PWMs are available for a TF, the most informative PWM was chosen, using the motif information score 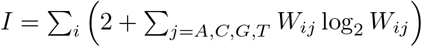 where *W_ij_* is the PWM probability for position *i* and nucleotide *j*. Also, similar motifs for distinct factors were identified with TOMTOM[21] and only the most informative was chosen. The selected motif PWM IDs are given in supplementary table S4).

In the section “Motif strength correlates with spatial interaction” we use the log likelihood ratio of a site as its “motif score”. Given a sequence *S* of length *L* and a PWM *W* of the same length, the 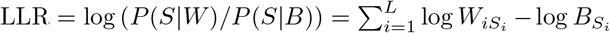 where *B* is a background model for the sequence (probability 0.25 per nucleotide).

## Results

### TFs fall in two broad groups

We explored co-binding of TFs in spatially proximal chromatin regions identified from ChIA-PET Pol II of GM12878 and K562 for 62 and 61 TFs respectively using their TFBS data from ChIP-seq peaks as described in Methods. For each pair of TFs, *p*-values were calculated for their co-occurrence based on 1000 randomizations. From this list of *p*-values, *q*-values were calculated using the Benjamini-Hochberg method[19]. The resulting matrix of *q*-values was clustered along both the rows and columns simultaneously for the GM12878 and K562 cell lines (figure 2).

**Figure 2:**
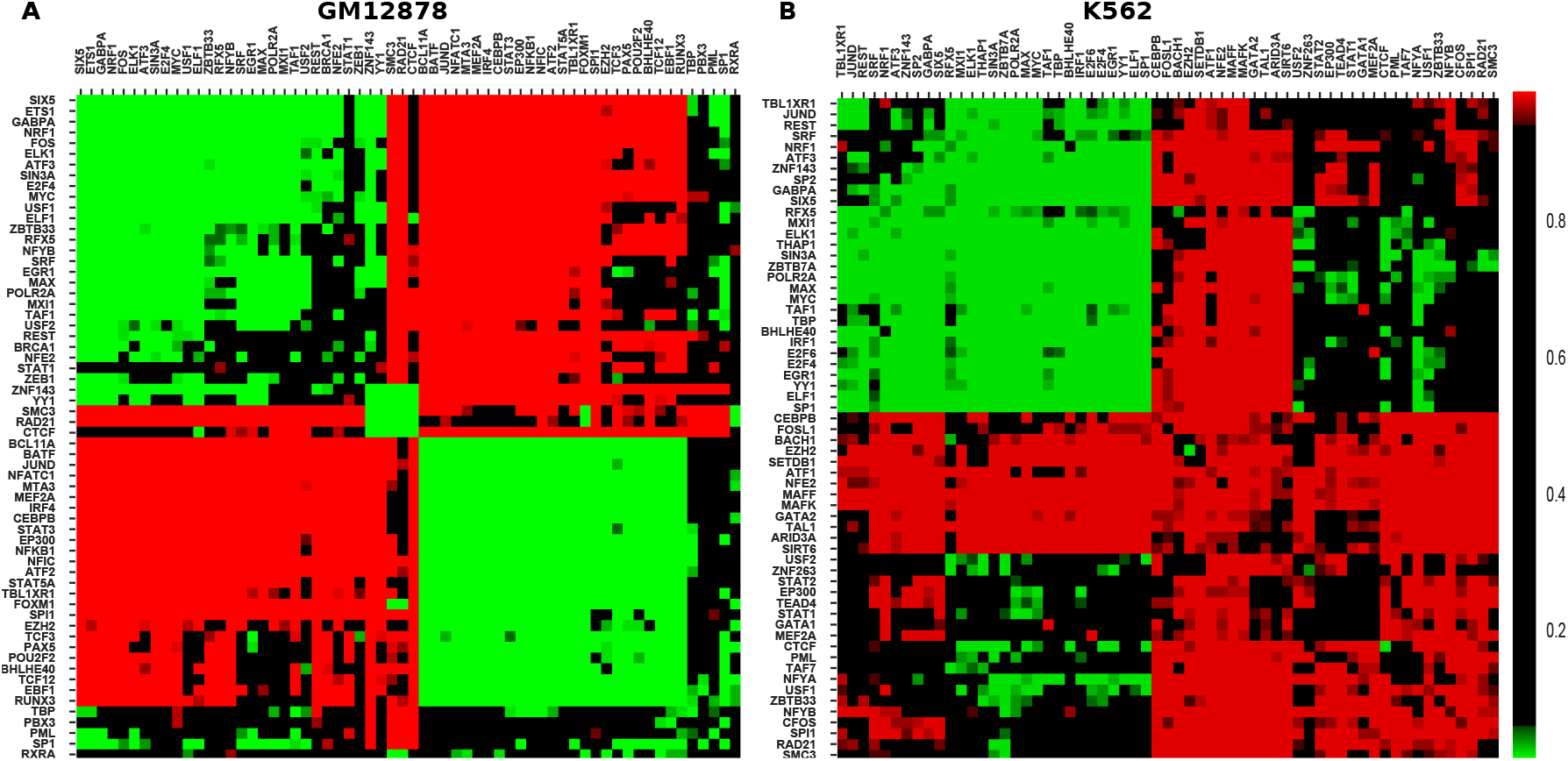
Clustered *q*-value heatmap showing attracting and repelling TF pairs in green and red respectively as described in Methods for (A) GM12878 and (B) K562 cell lines using ChIA-Pet polII data

In the GM12878 cell line, TFs segregate into two groups (Figure 2A). TFs in each group tend to attract other members of the same group (green, top left and bottom right of heatmap) but repel members of the other group (red, top-right, bottom-left). We call the top-right group “Group 1” and the bottom-right group “Group 2”.

A similar study, using a different methodology, was performed by Ma *et al*.[15]. Figure S2 in the supplementary material compares the results of the two studies; with a very few exceptions, our results are consistent with those of Ma et al.

In the K562 cell line, we do not observe two distinct classes of attracting TFs as in GM12878; instead, one attracting group emerges on the top-left, and most other TF pairs repel. However, as discussed further below, many of the characteristics of Group 1 from GM12878 are retained in the one attracting group in K562. We further discuss the differences between Group 1 and Group 2 in later subsections.

Notably, in both GM12878 and K562, CTCF and cohesin subunits SMC3 and RAD21 almost universally repel other TFs and mostly co-occur among themselves. Cohesin and CTCF have previously been reported to co-occur in spatially proximal regions and this co-occurrence is believed to be a critical for the formation of 3D chromatin loops[22, 23]. Pairs of CTCF motifs are found in a divergent orientation near cohesin, and the chromatin “loop extrusion model” has been proposed for chromatin organization via “topologically associated domains” (TADs). In this model, a loop of DNA is pushed through a cohesin ring, until it is hindered by CTCF molecules bound at the motif sites. This has been recently observed *in vitro* [24]. Our observations support the proposed interplay of cohesin and CTCF, and further suggest that the binding regions of cohesin in particular tend not to be in close spatial proximity with either promoters or other distal regulatory regions where other TFs tend to bind. Figure 3, in the next section, suggests that in addition to not being in spatial proximity of other TFBS, cohesin are also not in close sequential proximity of other TFBS.

**Figure 3:**
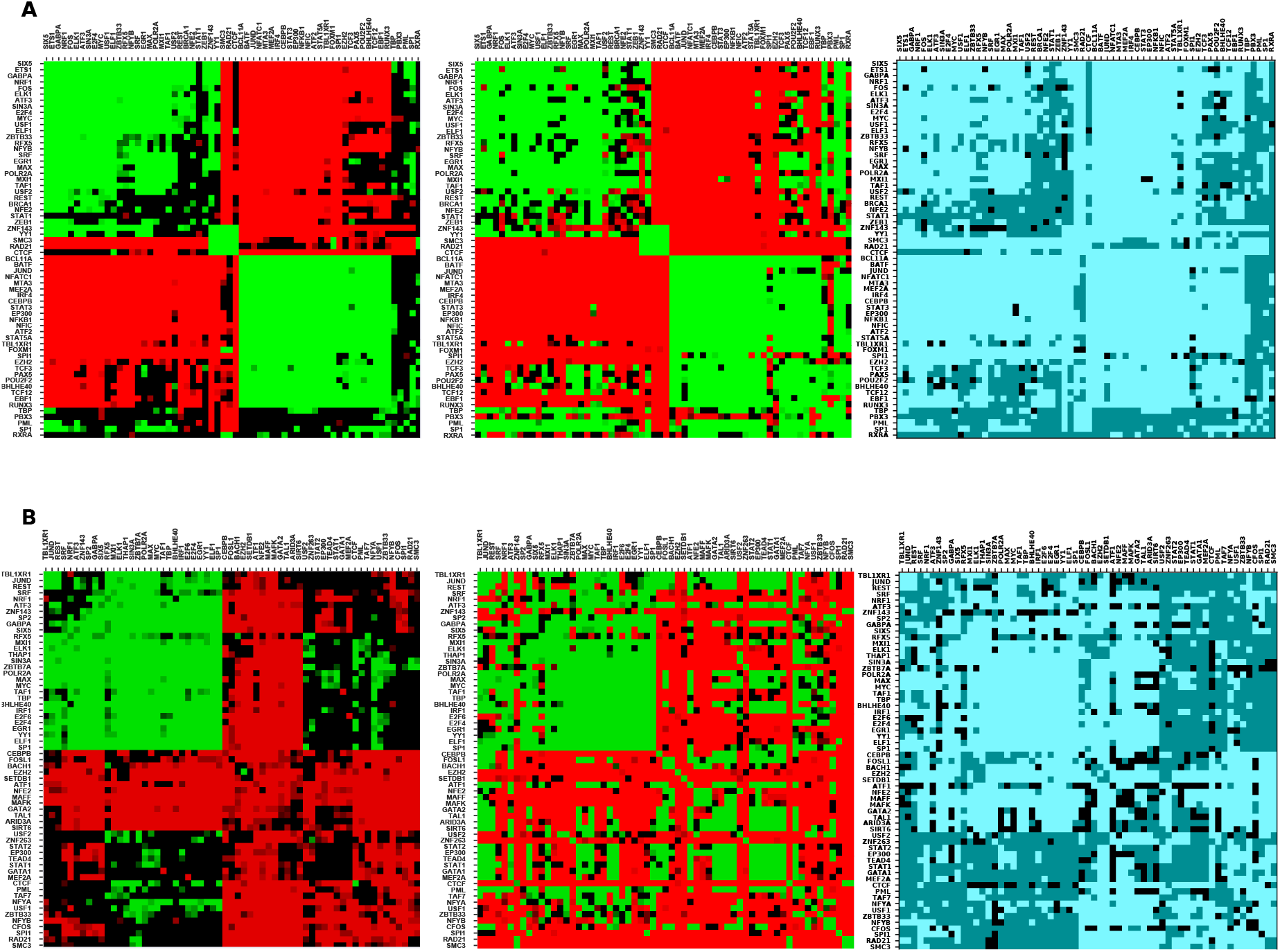
Top and bottom panels show the co-occurrence pattern in spatially proximal and sequentially contiguous regions for GM12878 and K562 cell lines respectively. The left heatmap in both panels corresponds to spatial co-occurrence, the middle to sequential co-occurrence, while the right heatmap is a comparison of these, coded as follows: both significant, in agreement: bright blue; both significant, in disagreement: black; one or both insignificant: dark blue.

In addition to TFs, we considered histone marks in our GM12878 analysis. The resulting attracting and repelling marks and TFs are shown in supplementary figure S3. We observe that histone marks associated with active promoters or enhancers i.e. H3K4me1, H3K4me2, H3K4me3, H3K27ac, and H3K9ac significantly co-occur among themselves even in spatially proximal regions similar to what has been reported in various previous studies based on sequentially contiguous regions of the genome [25, 26, 27, 28] and also co-occur with factors TBP, PML, and SP1 which are usually associated with the promoter regions. Also, these histone marks significantly repel their antagonistic marks H3K27me3 and H3K9me3.

Previous chromatin interaction studies have shown that the genome is primarily partitioned into distinct two compartments A and B, characterised by open and closed chromatin respectively. [4, 23]. It would be interesting to know if these compartments have any relation, or more importantly, if they contribute to the segregation of TFs that we observed in our investigation. To this end, we investigated the co-occurrence of TFs in the A and B compartments separately by classifying the spatial chromatin interactions falling into these compartments. But, we do not observe any such correlation of genome compartments with clusters of TFs in terms of their co-occurrence. The TF pair co-occurrence behaviour is given in supplementary figure S4A and S4B for A and B compartments respectively. The co-occurrence behaviour is similar in both compartments, but the numbers of significant attracting or avoiding TF pairs are fewer in the B compartment, as expected, it being generally inactive with a closed chromatin structure.

#### Sequential co-occurrence patterns largely mirror spatial co-occurrence

Prior to the advent of high-throughput chromatin interaction experiments such as Hi-C, co-occurrence of transcription factors was studied in sequentially contiguous regions such as promoters and enhancers [29, 30], and we asked whether the same patterns of attraction or repulsion occur within contiguous regulatory sequence. We repeated the analysis using the same regions from the ChIA-PET data, and the same randomization method, but this time considering co-occurrence within rather than across interacting regions. It turns out that most TF pairs exhibit the same qualitative behaviour as in spatially proximal regions; however, a few TF pairs show different qualitative behaviour (figure 3). These include interactions across groups: in particular, factors of Group 1 (SRF, EGR1, MAX, MXI1, TAF1) attract factors of Group 2 (TCF3, PAX5, POU2F2, BHLHE40, TCF12), visible as a green block in figure 3A (middle). Their pairwise spatial interactions are insignificant but they show significant co-occurrence in sequentially contiguous regions.

In GM12878, factors such as TBP, PML, SP1 which are usually enriched at the promoter regions attract almost all other TFs used in the study sequentially, but do not show any significant behaviour in spatially proximal regions, highlighting the promoter specific binding of these factors. In K562 cell line, as mentioned in the previous section, we observed only one main cluster of TFs attracting each other, but the other cluster of TFs, which do not attract spatially, do in many cases attract in sequentially contiguous regions. Overall, the analysis in spatial and sequential regions and comparison between them substantiate the evidence of two clusters of TFs present and suggest their potential role in the regulatory aspects.

#### Motif instances attract and repel similarly to TFBS

Our results above from the TF ChIP-seq binding data show considerable similarities and differences across cell lines in pattern of significant co-occurrence or avoidance in both spatially proximal and sequential contiguous regions. TFs in general bind to the genomic regions in a sequence specific manner and these short specific sequences are represented as sequence motifs. Motif instances of a given TF are present at several locations of genome, but at only at few of those sites does the TF bind in a given cell type depending on additional factors. We have so far considered binding events identified via ChIP-seq experiments. Here we ask whether a similar co-occurrence pattern occurs at a motif level, even though motif instances are generally poor indicators of tissue-specific TF binding.

We repeat the same analysis here, using motif occurrence rather than ChIP-seq binding information. Motif instances are predicted with FIMO[31] using motifs for TFs from the JASPAR database [20] as bundled with the MEME suite[32].

Motif information is available for a large number of TFs beyond the ones for which ChIP-seq data is available. We can study motif co-occurrence in various cell lines, while motif matches (unlike chip-seq peaks) are independent of cell lines, the chromatin contact information is cell line-dependent. We predict motif instances with FIMO[31] using motifs for TFs from the JASPAR database [20] as bundled with the MEME suite[32], specifically the file `JASPAR2018_CŨRE_vertebrates_redundant.meme` Rather than use all 719 motifs in that file, we identified similar motifs using TOMTOM[21] and clustered them into 93 groups, in each of which we picked the most informative motif as a representative (The selected TFs and their motif ids are given in supplementary material table S3).

Using these 93 distinct motifs, we observe strongly similar patterns of co-occurrence of motif sites in spatially proximal regions for four cell lines: GM12878(A), K562(B), MCF-7(C) and HeLa-S3(D), as shown in figure 4.

**Figure 4:**
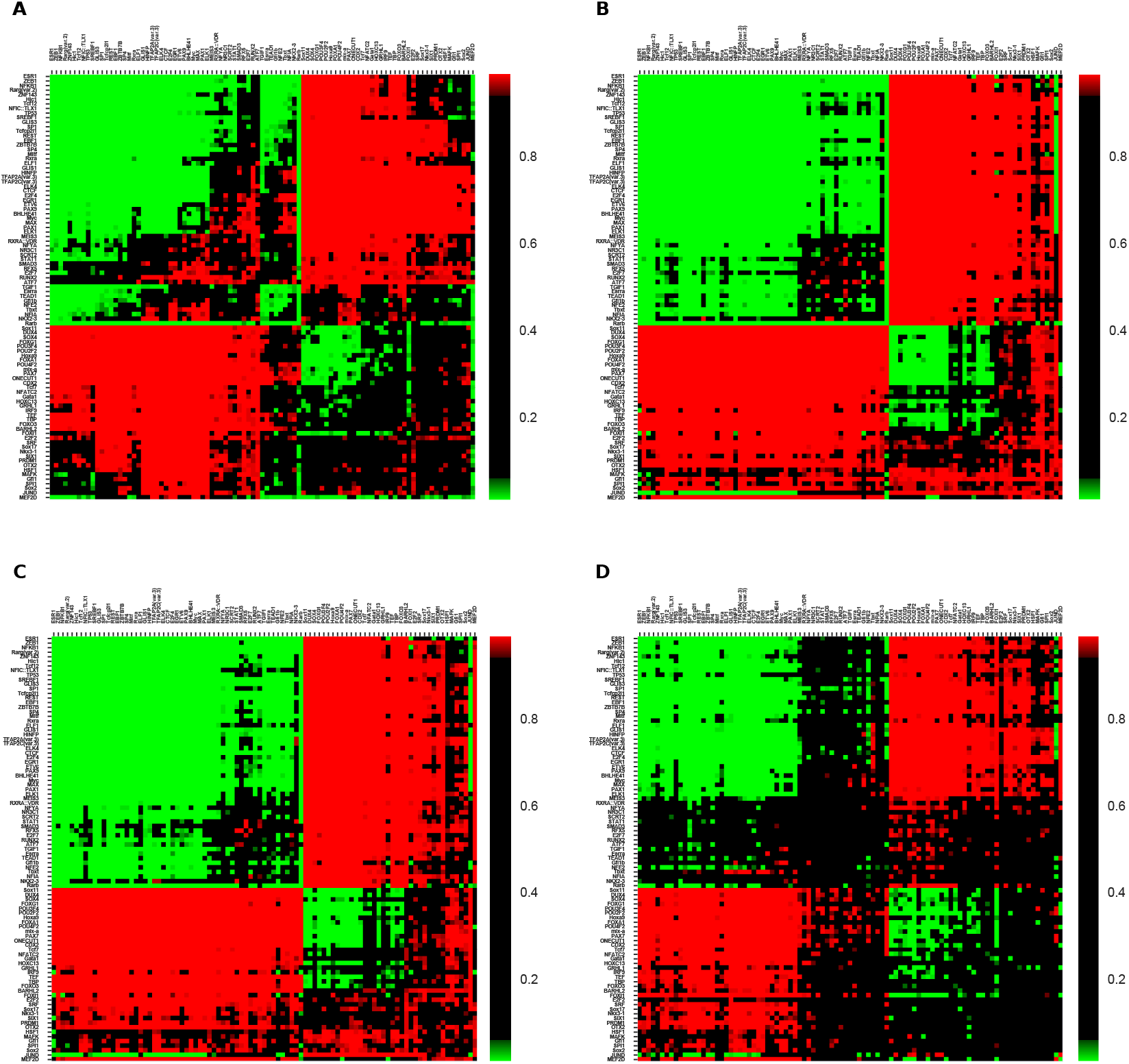
Comparison of motif co-occurrence in four cell lines: (A) GM12878, (B) K562, (C) MCF7 and (D) HeLa-S3. For ease of comparison, the order of factors from the clustering in GM12878 is used in all four subfigures

Motif co-occurrence in sequentially contiguous regions too is broadly similar to the spatially proximal regions, in all the four cell lines examined. This can seen in the supplementary figures S7, S8, S10 and S9 for GM12878, K562, MCF7 and HeLa-S3 cell lines respectively. However, there are many more significant examples both of attraction and of repulsion in sequentially contiguous regions, suggesting perhaps a greater degree of combinatorial control within promoters and enhancers.

#### A consensus TF-TF co-occurrence network

Using the co-occurrence patterns for binding events and motifs that we see in spatially and sequentially contiguous regions, we propose a consensus network for TF pair co-occurrence. We use both motif and experimental (ChIP-seq) binding events, and both spatially proximal and sequentially contiguous regions, giving us four datasets; and we require that TF pairs attract in at least two of these four datasets. The consensus network is shown in figure 5A and 5B for GM12878 and K562 cell lines respectively. A comparison of observations from all four types is shown in supplementary figures S11 and S12. This high confidence consensus network further highlights the segregation of TFs into two groups with certain TFs such as TBP, PBX3, SP1, REST, TCF12, TCF3, and PAX5, acting as intermediates, while cohesin subunits SMC3 and RAD21 are outliers.

**Figure 5:**
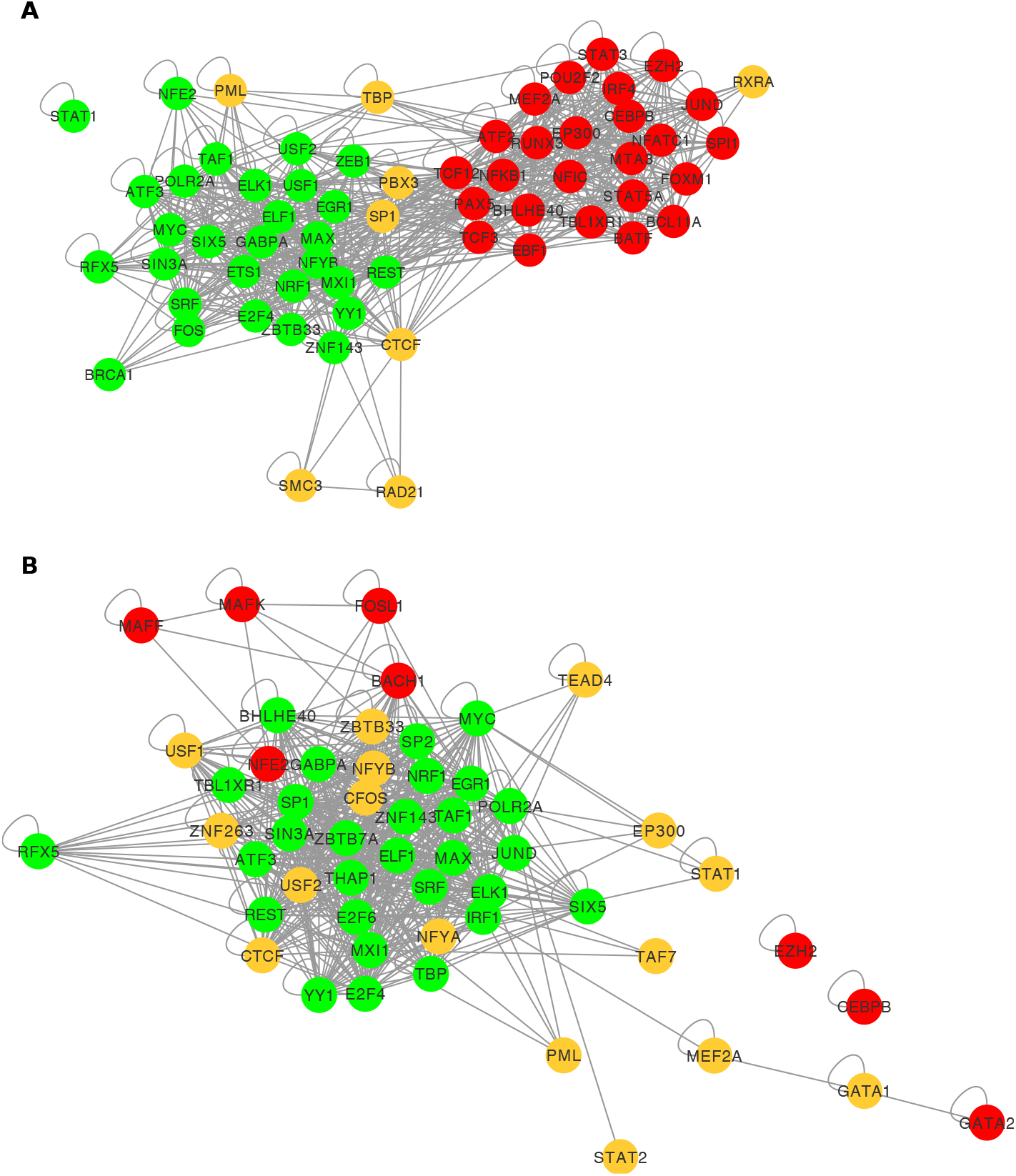
High confidence consensus networks built using the TF pair co-occurrence observations from all four methods (spatial or sequential, binding or motif instance) for (A) GM12878 and (B) K562 cell lines.

### The two main TF groups interact differently with proteins, DNA, and genes

In GM12878 TFs separate into two groups that attract withing a group, but repel across groups, with some exceptions. In K562 there is one group that mutually attracts (and largely mirrors Group 1 in GM12878), and another that repels most other TFs in our dataset. The two groups in each case exhibit differences in their interactions with other proteins, DNA binding locations, and downstream gene targets.

#### PPI interactions are enriched in intra-group TF pairs in GM12878

We examined physical interactions of TF-TF pairs using the Human Integrated Protein-Protein Interaction Reference (v2.2) database (HIPPIE) [33]. We observe enrichment of physical interaction among attracting TF pairs (92 out of 539 i.e. ~0.17) compared to avoiding TF pairs (40 out of 523 i.e. ^0.07) indicating that attracting TF pairs are significantly more likely to interact physically than avoiding pairs (*p* = 1.98 × 10^-6^, hypergeometric test). There is no significant difference between Group 1 TF pairs and Group 2 TF pairs (interacting fraction ~0.15 and ~0.18 respectively).

#### Domain-domain interactions are enriched in attracting TF pairs

To further examine the possibility of undocumented protein-protein interactions, we considered the domain structures of the TFs and looked for possible domain-domain interactions among all TF pairs present in the study, using the database of three-dimensional interacting domains (3did) [34]. Any pair of TFs containing respective interacting domains, was considered “potentially interacting”. Attracting TF pairs in both cell lines show significant enrichment over avoiding TF pairs for potential physical interaction (tables 1 and 2).

**Table 1:** The number of possible physical interactions from the literature of domain-domain physical interactions among attracting and avoiding TF pairs for GM12878 cell line. Attracting pairs are significantly more likely to have possible domain-domain interactions (*p* = 2.17 × 10^-9^, hypergeometric test)

|  | Attracting TF pairs | Avoiding TF pairs | Total |
| --- | --- | --- | --- |
| potential physical interactions | 131 | 61 | 192 |
| No potential physical interactions | 598 | 718 | 1316 |
| Total | 729 | 779 | 1508 |

**Table 2:**
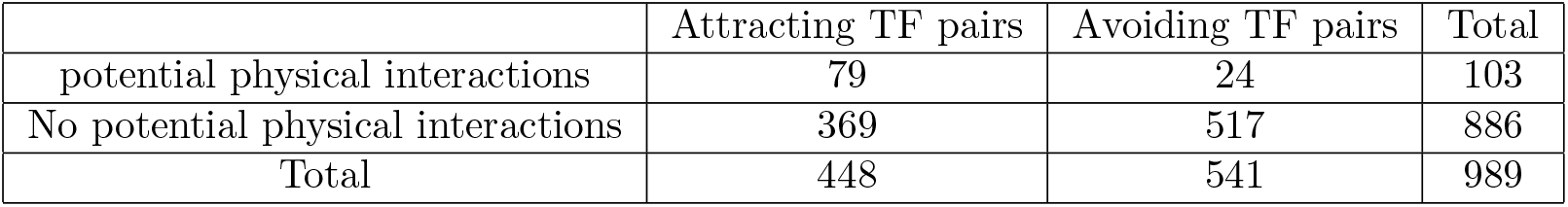
Similar to table 1, for K562 cell line. Attracting pairs are significantly more likely to have possible domain-domain interactions (*p* = 7.65 × 10^-12^, hypergeometric test)

#### Internal nodes in PPI pathways largely differ in Group 1 and Group 2 for GM12878

We next looked at TFs that may be binding indirectly, via co-factors. We considered the shortest PPI pathway for each pair of TFs. Specifically, we asked what TFs (not necessarily in our list of 62) occur in the shortest interaction pathway between that pair of TFs, and evaluated the frequency of occurrence of each such “internal node”. We only considered annotated TFs, and not other proteins (figure 6). Interestingly, the frequent internal nodes are largely unique for each group. Moreover, the few TFs common to these groups tend to occur as internal nodes of internal-group TF pairs, suggesting possible roles as co-activators or co-repressors. Taken together, this analysis suggests that the two groups have distinct structural underpinnings.

**Figure 6:**
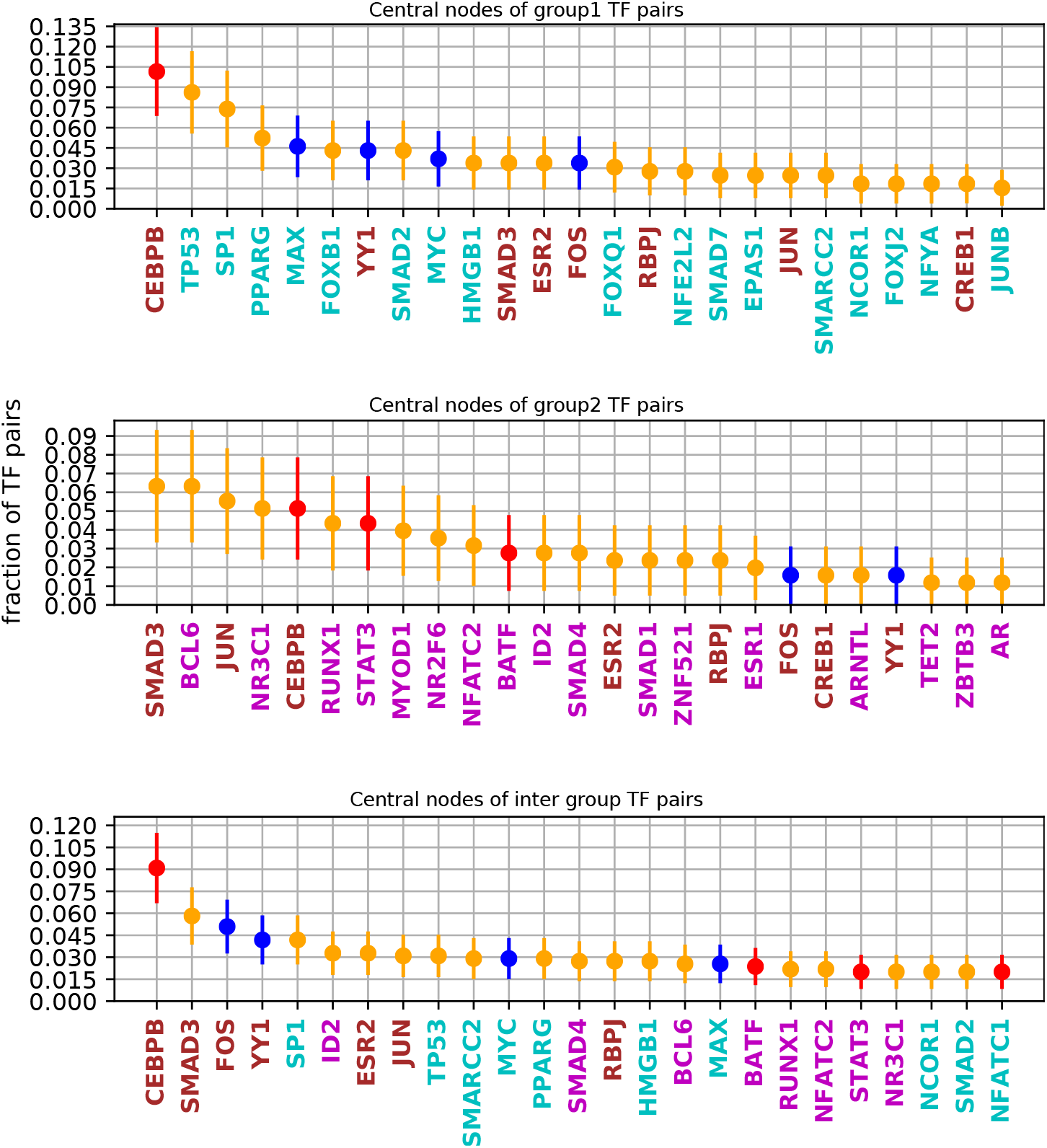
The TFs most frequently occurring as internal nodes in the shortest PPI pathway between TF pairs, for pairs from Group 1 (top), Group 2 (middle), and cross-group(bottom panel). The blue, red markers are Group 1, Group 2 TFs respectively and yellow are TFs not present in our co-occurrence data. TFs in cyan text occur as central nodes within Group 1 only, in magenta are central nodes within Group 2 only, and brown occur as central nodes in both groups.

#### Group 1 TFs bind closer to promoters than Group 2 TFs

Figure 7 plots the cumulative distribution of ChIP-seq peak distance from the nearest transciption start site (TSS) for Group 1, Group 2 and ungrouped TFs, in (A) GM12878 and (b) K562. Group 1 TFs tend to bind closer to the TSS than Group 2 or ungrouped TFs. This is consistent with what Ma *et al*[15] report.

**Figure 7:**
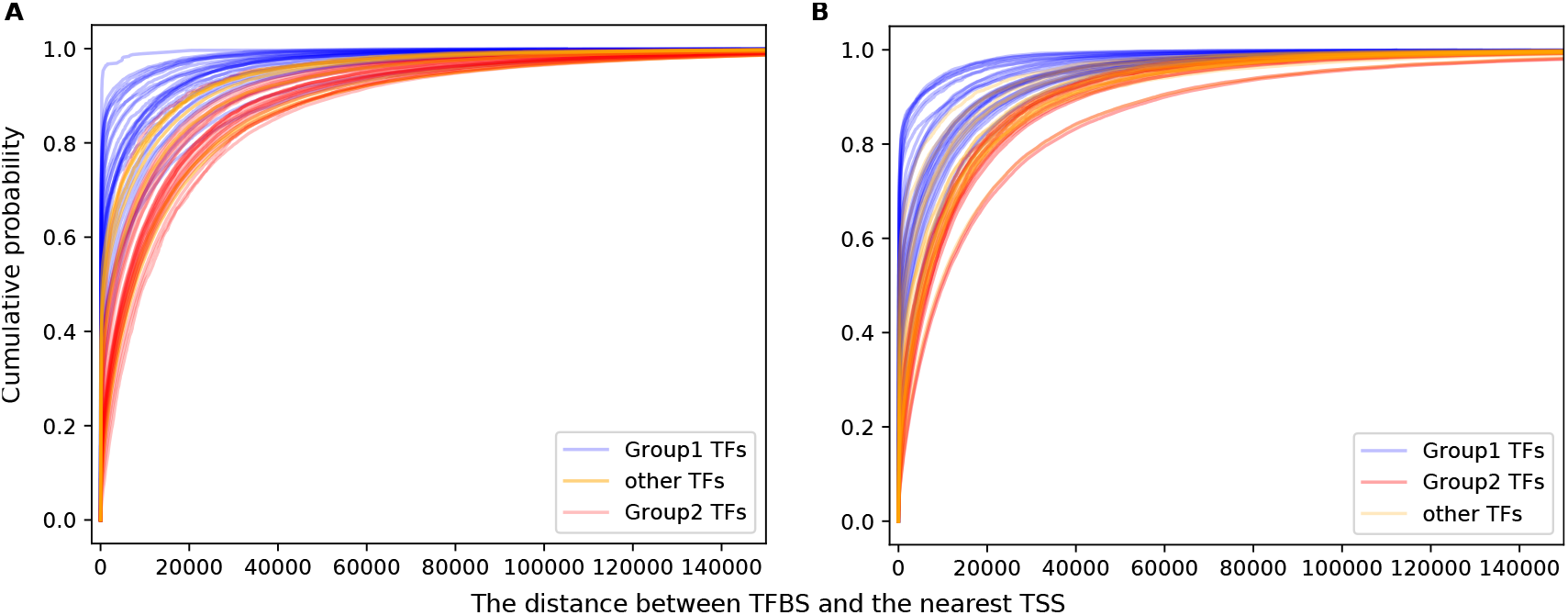
(A) The distance between TFBS and the nearest TSS, and the cumulative probability for sites to occur within that distance, for Group 1 TFs (blue), Group 2 TFs (red) and other TFs (yellow), in (A) GM12878 and (B) K562.

#### Group 1 target genes are enriched for housekeeping functions, Group 2 for tissue-specific functions

We identify putative target genes of each TF as genes located either within 2kbp sequentially of a TFBS for that factor, or on a spatially proximal region to that TFBS. Within each group, We consider the functions of target genes, using Gene Ontology (GO) functional enrichment analysis and disease trait enrichment analysis.

Based on the distributions of putative TF regulators per gene (supplementary figure S14), genes that are targets for at least five TFs from one group, and at most two TFs from the other group, are taken to be target genes for the former group. We perform GO term enrichment analysis for biological processes, and molecular functions using DAVID [35] with background of all human genes. We find that Group 1 target genes are enriched for housekeeping functions such as transcription, cell cycle, transport, and metabolism, while Group 2 target genes are enriched for immune/inflammatory response (figure 8A).

**Figure 8:**
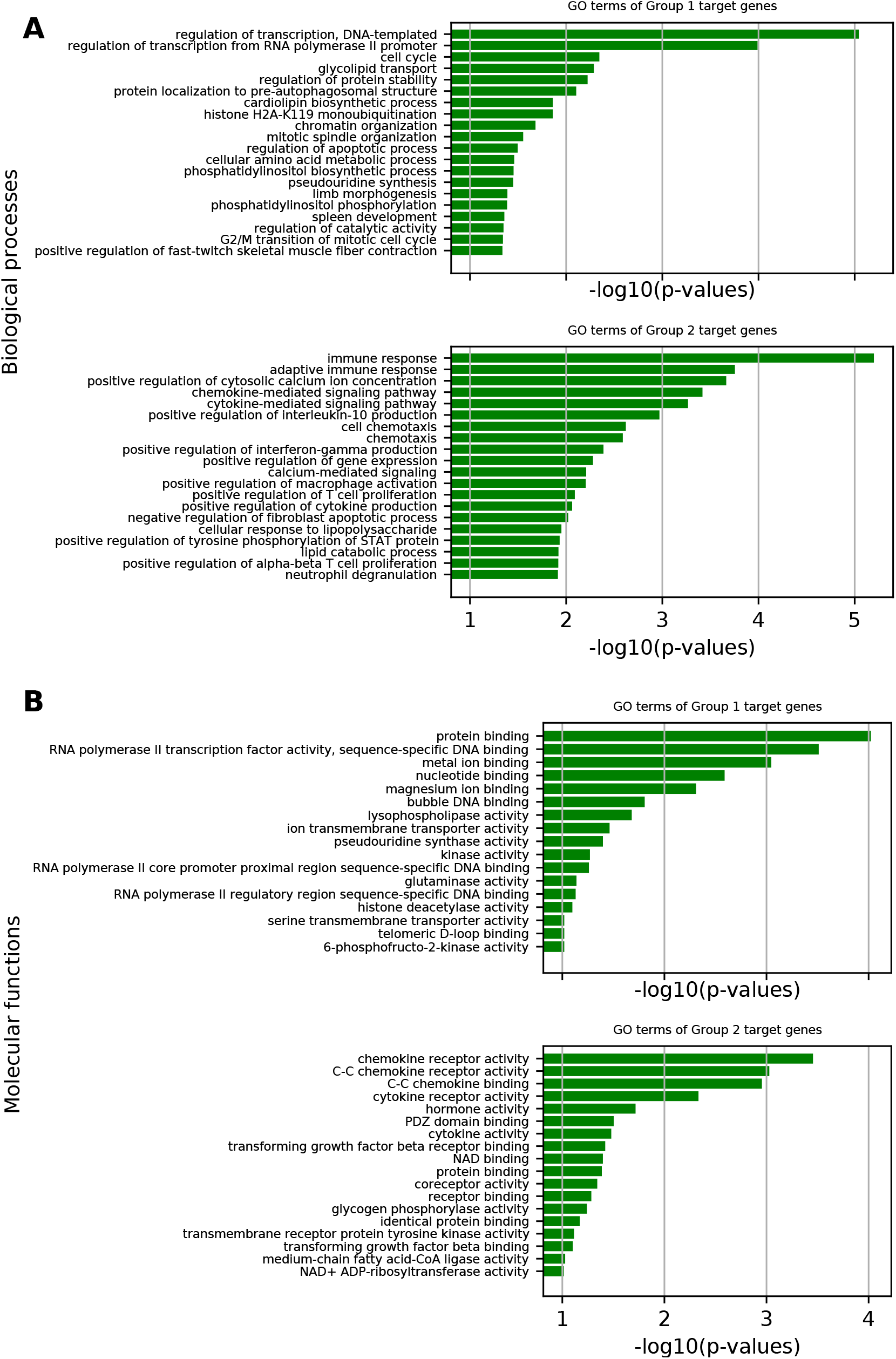
(A) shows the enriched biological process GO terms for the Group 1 and Group 2 target genes. (B) Similarly, enriched GO molecular function terms are shown for Group 1 and Group 2 target genes.

We specifically assessed whether Group 1 target genes are enriched for housekeeping genes using a database of known housekeeping genes [36]. Group 1 target genes are significantly enriched over Group 2 target genes for housekeeping genes (hypergeometric test *p*-value 2.7 × 10^-9^). The enrichment is not highly sensitive to the target gene selection criteria (for example, requiring 3 TFs from one group and 0 from the other yields similar results).

The biological process enrichment is corroborated by molecular function enrichment. The molecular functions of Group 1 target gene involve mostly DNA binding, metal ion binding events, while Group 2 genes involves in receptor activity of chemokines and cytokines, involved in immune responses (figure 8B).

Next, we assessed the group-specific target genes for association with disease traits from the GWAS catalog [37], only considering the traits associated with at least 5 genes. The target genes of each group were examined for significant overlap with these traits (Fisher exact test *p* < 0.05). Consistently, Group 2 targets were enriched for inflammatory disease, immunological diseases involving lymphocyte, leukocytes, and neutrophils, while Group 1 genes are not particularly enriched for any particular diseases (supplementary figure S16). Enrichment of immune response and traits in Group 2 targets is especially interesting considering that GM12878 is a lymphocytic cell line.

We performed a similar analysis for K562 cell line treating the large group of mutually attracting TFs as Group 1 and the remainder as Group 2. Again, Group 1 target genes are enriched for housekeeping functions such as transcription, metabolism, and transport (hypergeometric test *p*-value 1.4 × 10^-3^). Group 2 target genes are not particularly enriched for pathway or functions (supplementary figure S17).

Finally, in the HeLa-S3 cell line we see a similar segregation into two groups as in GM12878, but Group 1 is much larger than Group 2 (figure S5). Again, Group 1 is significantly enriched for housekeeping functions (*p* = 4.9 × 10^-17^) (figure S18). Group 2 is enriched for protein ubiquitination and various metabolic processes, but with little overlap to Group 2 of GM12878.

Both the similarity and differences across the cell lines in terms of TF groupings and their target functions are worth noting. GM12878 is a lymphoblastoid cell line derived from the blood of a healthy female donor, K562 is lymphoblasts isolated from the bone marrow of a chronic myelogenous leukemia patient, and HeLa-S3 is a cervix carcinoma cell line. While group-1 TF targets in all three cell lines are enriched for house keeping functions, Group 2 of TFs is most prominent in GM12878, derived from a healthy donor, and their targets are enriched for immune function. In contrast, leukemia-derived K562 lacks a functionally coherent Group 2 TFs, which may suggest a loss of lineage-specification in leukemic lymphoblasts. HeLa-S3 has a smaller Group 2 than GM12878 with a divergent set of, potentially lineagespecific, target functions.

### Motif strength correlates with spatial interaction

Previous studies have demonstrated the significance of long-range spatially proximal regions in aiding *in vivo* TF binding at weaker motif sites by containing several homotypic motif sites in spatially clustered genomic regions [14, 15]. We extend the investigation to spatial interactions with other genomic regions, regardless of whether there are TFs binding to those other regions.

For each TF we consider the most informative PWM available in JASPAR[20] (see Methods). Individual binding sites may have stronger or weaker matches to the PWM, which is an estimate of their binding affinity to the TF. We use the log likelihood ratio (see Methods) as the “motif score” for a site, a measure of the likely “strength” of binding at that site. For each TF, we consider peaks containing sites whose motif scores are among among the top 10% as “peaks with strong motifs”. We then compare the fraction of peaks with strong motifs between two groups of peaks—those that are isolated (occurring in regions without any spatial chromatin interactions) and those in spatially interacting regions.

We observe for almost all factors, spatially isolated TFBS exhibit a significantly greater fraction of strong motifs (Figure 9A). If we consider whether the spatially interacting region is bound by the same TF or is not (but possibly bound by another TF), in the majority of cases binding by the same TF is associated with a weaker motif, but this is highly variable (Figure 9A). A similar trend is observed in K562 cell line (supplementary figure S13). This suggests that spatial interactions enable binding by TFs to weaker motifs, even in the absence of homotypic binding or the presence of heterotypic binding as was suggested previously.

**Figure 9:**
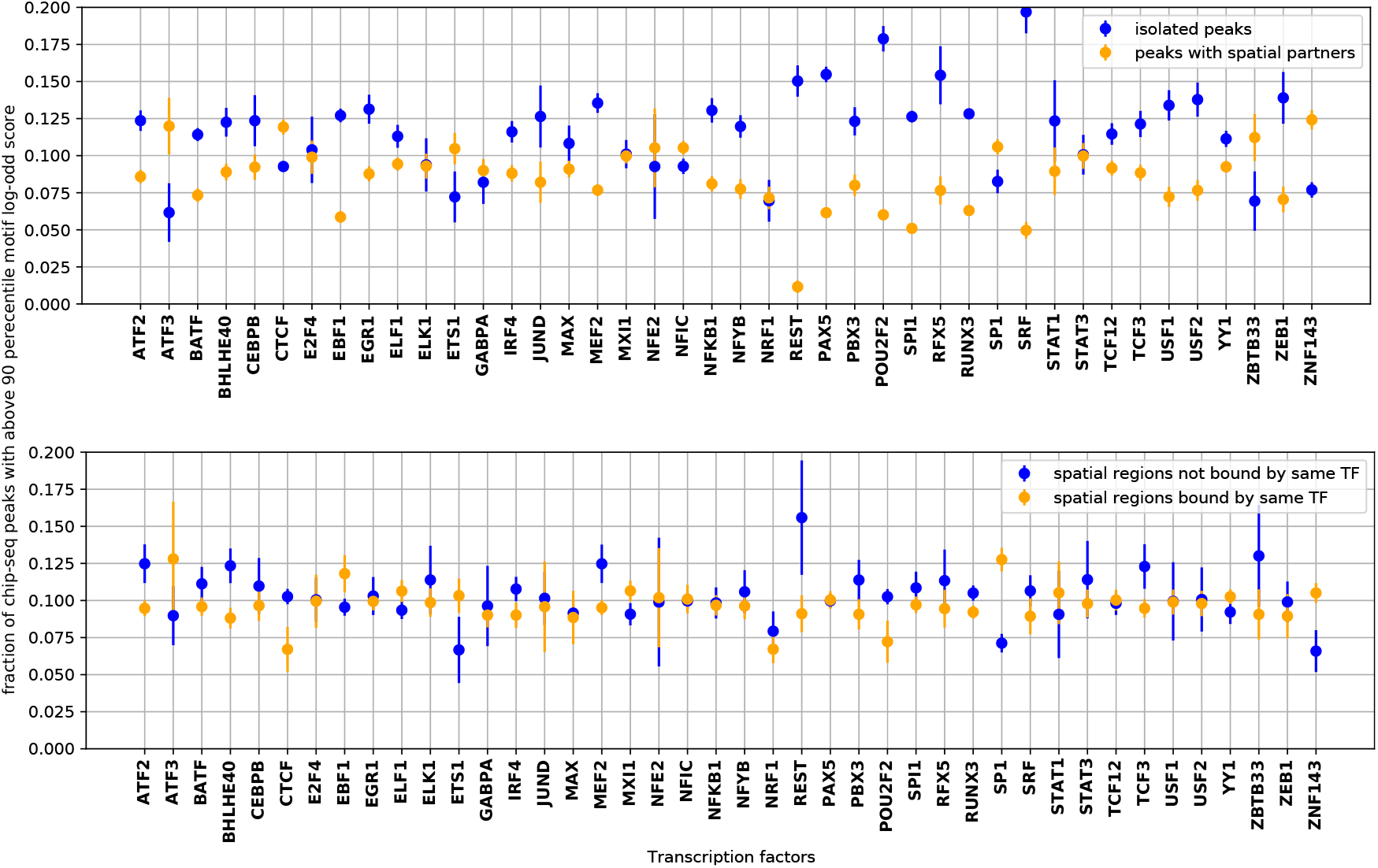
Spatially isolated ChIP-seq peaks mostly have a higher fraction of strong motifs than peaks within spatial clusters (top panel). For peaks within spatial clusters, when the same TF binds to a spatially-interacting region, the associated motif is weaker in 24 out of 42 cases (bottom panel).

## Discussion

We present a methodology to assess significant co-occurrence, and significant avoidance, between pairs of transcription factors in both linear as well as spatially proximal regions in genome. We find that most TF pairs attract or repel (FDR *q* ≤ 0.05) in every cell type that we examined. The cohesin subunits RAD21 and SMC3 tend to repel most other factors across cell lines. CTCF tends to repel many factors, but attracts others. This suggests that regions where cohesin rings form are not transcriptionally active [38, 39, 40, 41, 42], while CTCF, which plays multiple roles [43] including chromatin organization, insulator function, and transcriptional regulation, can attract or avoid other factors based on function. A few factors, notably promoter-associated factors such as SP1 and TBP, co-occur widely with factors in both groups.

TF co-occurrence pattern is corroborated by co-occurrence of histone marks in GM12878: marks associated with active promoters/enhancers co-occur with one another and also with TBP, PML and SP1 and a large group of TFs, while repelling a smaller group of other histone marks, including marks associated with heterochromatin. Heterochromatin marks repel other histone marks and most TFs. In GM12878, we also find substantial agreement with a previous analysis based on a different methodology [15].

Sequentially proximal binding shows largely the same pattern of co-occurrence as the spatially proximal binding but with greater significance. Furthermore, motif instances too show co-occurrence similar to ChIP-seq binding events, overall suggesting a potentially coordinated evolution of cis elements with the chromatin structure. We synthesise the various co-occurrence data (spatial and sequential; TF binding and motif instance) into a consensus co-occurrence network that illustrates these points (figure 5A).

We find that the attraction-repulsion patterns and the separation into groups is reflected in various biological properties of the TF pairs and groups, including proximity from TSS, protein interactions networks, domain-domain interactions, and function of downstream target genes. These too exhibit similar trends across cell lines, with some differences. The similarities suggest that basic transcriptional machinery depends on the chromatin structure bringing appropriate regions together; while the differences that do occur suggest that chromatin state plays an important function in cell-type differentiation. Specific differences in the revealed TF groups among K562, HeLa-S3 and GM12828 suggests that while Group 1 TFs associated with housekeeping functions are conserved between the three cell lines, the Group 2 TFs associated with lineage-specific functions are disrupted specifically in the malignant lymphoblast-derived K562 and cervical carcinoma-derived HeLa-S3.

Finally our method provides a platform for testing heterotypic as well as homotypic spatial interactions of TFs. While previous studies have suggested the role of homotypic binding in spatially clustered regions in boosting *in vivo* binding [14], our analysis shows that proximity to other regulatory regions is associated with stronger in vivo binding, regardless of TF binding in the spatially proximal regions.

Overall, our work indicates a complex interplay of TF binding and chromatin structure, some of which we elucidate in detail, while much remains to be discovered in future. Our tool, ChromTogether, will facilitate future studies in this field.

## Acknowledgements

Ankit Agrawal helped with early data collection and analysis.

## Author contributions

SH, LN, RS conceived the early outlines of the work. All authors contributed to further development of the work. RNV performed all data collection, software development, and analysis. All authors contributed to writing the manuscript. All authors have read and approved the final manuscript.

