## Supplementary Information for "Transcription factors organize into functional groups on the linear genome and in 3D chromatin"

#### Supplementary figures and tables

#### Materials & Methods

| Cell line | Data Set | Accession Number |
|---|---|---|
| GM12878 | ChIA-PET (RNAPII) | GSM1872887 |
| K562 | ChIA-PET (RNAPII) | GSM970213 |
| MCF7 | ChIA-PET (RNAPII) | GSM970209 |
| HeLa-S3 | ChIA-PET (RNAPII) | GSM1872889 |

Table S1: Accession IDs of chromatin interaction data sets used in the study

| **GM12878** | |
|---|---|
| ATF2 | wgEncodeEH002306 |
| ATF3 | wgEncodeEH001562 |
| BATF | wgEncodeEH001479 |
| BCL11A | wgEncodeEH001486 |
| BHLHE40 | wgEncodeEH002025 |
| BRCA1 | wgEncodeEH001830 |
| CEBPB | wgEncodeEH003212 |
| CTCF | wgEncodeEH001851, wgEncodeEH000029, wgEncodeEH000394, wgEncodeEH000532 |
| E2F4 | wgEncodeEH002867 |
| EBF1 | wgEncodeEH001832 |
| EGR1 | wgEncodeEH002328 |
| ELF1 | wgEncodeEH001617 |
| ELK1 | wgEncodeEH002851 |
| EP300 | wgEncodeEH002824, wgEncodeEH002037 |
| ETS1 | wgEncodeEH001564 |
| EZH2 | wgEncodeEH002411 |
| FOS | wgEncodeEH000622 |
| FOXM1 | wgEncodeEH002529 |
| GABPA | wgEncodeEH001462 |
| IRF4 | wgEncodeEH001484 |
| JUND | wgEncodeEH000639 |
| MAX | wgEncodeEH002806 |
| MAZ | wgEncodeEH002852 |
| MEF2A | wgEncodeEH001565 |
| MTA3 | wgEncodeEH002329 |
| MXI1 | wgEncodeEH002026 |
| MYC | wgEncodeEH000547 |
| NFATC1 | wgEncodeEH002307 |
| NFE2 | wgEncodeEH001808 |
| NFIC | wgEncodeEH002343 |
| NFYB | wgEncodeEH002065 |
| NRF1 | wgEncodeEH001846 |
| PAX5 | wgEncodeEH001489, wgEncodeEH001495 |
| PBX3 | wgEncodeEH001477 |
| PML | wgEncodeEH002308 |
| POLR2A | wgEncodeEH000626, wgEncodeEH001517 |
| POU2F2 | wgEncodeEH001475 |
| RAD21 | wgEncodeEH000749 |
| REST | wgEncodeEH002314 |
| RFX5 | wgEncodeEH001810 |
| RUNX3 | wgEncodeEH002330 |
| RXRA | wgEncodeEH001541 |

| | |
|---|---|
| SIN3A | wgEncodeEH002868 |
| SIX5 | wgEncodeEH001542 |
| SMC3 | wgEncodeEH001833 |
| SP1 | wgEncodeEH001496 |
| SPI1 | wgEncodeEH001476 |
| SRF | wgEncodeEH001464 |
| STAT1 | wgEncodeEH001852 |
| STAT3 | wgEncodeEH001811 |
| STAT5A | wgEncodeEH002321 |
| TAF1 | wgEncodeEH001478 |
| TBL1XR1 | wgEncodeEH002853 |
| TBP | wgEncodeEH001798 |
| TCF12 | wgEncodeEH001485 |
| TCF3 | wgEncodeEH002315 |
| USF1 | wgEncodeEH001468 |
| USF2 | wgEncodeEH001812 |
| YY1 | wgEncodeEH000695, wgEncodeEH001657 |
| ZBTB33 | wgEncodeEH001488 |
| ZEB1 | wgEncodeEH001645 |
| ZNF143 | wgEncodeEH001853 |
| **K562** | |
| ARID3A | wgEncodeEH002861 |
| ATF1 | wgEncodeEH002865 |
| ATF3 | wgEncodeEH000700 |
| BACH1 | wgEncodeEH002846 |
| BHLHE40 | wgEncodeEH001857 |
| CEBPB | wgEncodeEH001821 |
| CTCF | wgEncodeEH002279, wgEncodeEH000042 |
| CTCF | wgEncodeEH002797 |
| E2F4 | wgEncodeEH000671 |
| E2F6 | wgEncodeEH000676, wgEncodeEH001598 |
| EGR1 | wgEncodeEH001646 |
| ELF1 | wgEncodeEH001619 |
| ELK1 | wgEncodeEH003356 |
| EP300 | wgEncodeEH002834, wgEncodeEH002086 |
| EZH2 | wgEncodeEH002089 |
| FOS | wgEncodeEH001207, wgEncodeEH000619 |
| FOSL1 | wgEncodeEH001637 |
| GABPA | wgEncodeEH001604 |
| GATA1 | wgEncodeEH000638 |
| GATA2 | wgEncodeEH000683 |
| GATA2 | wgEncodeEH001208, wgEncodeEH001576 |
| IRF1 | wgEncodeEH001866, wgEncodeEH002798, wgEncodeEH002799 |
| JUND | wgEncodeEH002164 |
| MAFF | wgEncodeEH002804 |
| MAFK | wgEncodeEH001844 |
| MAX | wgEncodeEH002869 |
| MEF2A | wgEncodeEH001663 |
| MXI1 | wgEncodeEH001827 |
| MYC | wgEncodeEH002800, wgEncodeEH001867 |
| NFE2 | wgEncodeEH000624 |
| NFYA | wgEncodeEH002021 |
| NFYB | wgEncodeEH002024 |
| NRF1 | wgEncodeEH001796 |
| PML | wgEncodeEH002320 |
| POLR2A | wgEncodeEH000704, wgEncodeEH000727 |
| RAD21 | wgEncodeEH001585, wgEncodeEH000649 |
| REST | wgEncodeEH001638 |
| RFX5 | wgEncodeEH002033 |

| | |
|---|---|
| SETDB1 | wgEncodeEH000677 |
| SIN3AK20 | wgEncodeEH001607 |
| SIRT6 | wgEncodeEH000681 |
| SIX5 | wgEncodeEH001483 |
| SMC3 | wgEncodeEH001845 |
| SP1 | wgEncodeEH001578 |
| SP2 | wgEncodeEH001653 |
| SPI1 | wgEncodeEH001482 |
| SRF | wgEncodeEH001600 |
| STAT1 | wgEncodeEH000760, wgEncodeEH000761 |
| STAT2 | wgEncodeEH000665 |
| STAT5A | wgEncodeEH002347 |
| TAF1 | wgEncodeEH001582 |
| TAF7 | wgEncodeEH001654 |
| TAL1 | wgEncodeEH001824 |
| TBL1XR1 | wgEncodeEH002848, wgEncodeEH002849 |
| TBP | wgEncodeEH001825 |
| TEAD4 | wgEncodeEH002333 |
| THAP1 | wgEncodeEH001655 |
| USF1 | wgEncodeEH001583 |
| USF2 | wgEncodeEH001797 |
| YY1 | wgEncodeEH000684, wgEncodeEH001584, wgEncodeEH001623 |
| ZBTB33 | wgEncodeEH001569 |
| ZBTB7A | wgEncodeEH001620 |
| ZNF143 | wgEncodeEH002030 |
| ZNF263 | wgEncodeEH000630 |
| **HeLa-S3** | |
| BRCA1 | wgEncodeEH001814 |
| BRF1 | wgEncodeEH000764 |
| BRF2 | wgEncodeEH000765 |
| CEBPB | wgEncodeEH001815 |
| CHD2 | wgEncodeEH002027 |
| CTCF | wgEncodeEH000398, wgEncodeEH000541 |
| E2F1 | wgEncodeEH000688 |
| E2F1 | wgEncodeEH000699 |
| E2F4 | wgEncodeEH000689 |
| E2F6 | wgEncodeEH000692 |
| ELK1 | wgEncodeEH002864 |
| ELK4 | wgEncodeEH001753 |
| EP300 | wgEncodeEH001820 |
| EZH2 | wgEncodeEH003086 |
| FAM48A | wgEncodeEH001855 |
| FOS | wgEncodeEH000647 |
| GABPA | wgEncodeEH001504 |
| GTF2F1 | wgEncodeEH001816 |
| IRF3 | wgEncodeEH001788 |
| JUN | wgEncodeEH000746 |
| JUND | wgEncodeEH000745 |
| MAFK | wgEncodeEH002856 |
| MAX | wgEncodeEH002830 |
| MAZ | wgEncodeEH002855 |
| MXI1 | wgEncodeEH001826 |
| MYC | wgEncodeEH000542, wgEncodeEH000648 |
| NFYA | wgEncodeEH002066 |
| NFYB | wgEncodeEH002067 |
| NR2C2 | wgEncodeEH000687 |
| NRF1 | wgEncodeEH000723 |
| POLR2A | wgEncodeEH001474, wgEncodeEH001838 |
| PRDM1 | wgEncodeEH001817 |

| | |
|---|---|
| RAD21 | wgEncodeEH001789 |
| RCOR1 | wgEncodeEH002844 |
| REST | wgEncodeEH001629 |
| RFX5 | wgEncodeEH001818 |
| RPC155 | wgEncodeEH000766 |
| SMC3 | wgEncodeEH001839 |
| STAT1 | wgEncodeEH000614 |
| STAT3 | wgEncodeEH001799 |
| TAF1 | wgEncodeEH001505 |
| TBP | wgEncodeEH001790 |
| TCF7L2 | wgEncodeEH002069, wgEncodeEH002813 |
| USF2 | wgEncodeEH001819 |
| ZKSCAN1 | wgEncodeEH002857 |
| ZNF143 | wgEncodeEH002028 |
| ZNF274 | wgEncodeEH001763 |
| ZZZ3 | wgEncodeEH001872 |

Table S2: Accession IDs of the uniformly processed chip-seq peaks generated by ENCODE project used in the study

The commands for various external tools that were used in the study are given below.

For scanning presence of motif instances in chromatin regions, FIMO[31] was used with following command:

```
fimo -o <output-directory> <TF-motif-meme-file> <fasta-sequence-file-of-chromatin-regions>
```

For finding the similarity between all the nonredundant JASPAR motifs, we used the TOMTOM[21] external tool:

```
tomtom <query-motif-file> <target-motif-file>
```

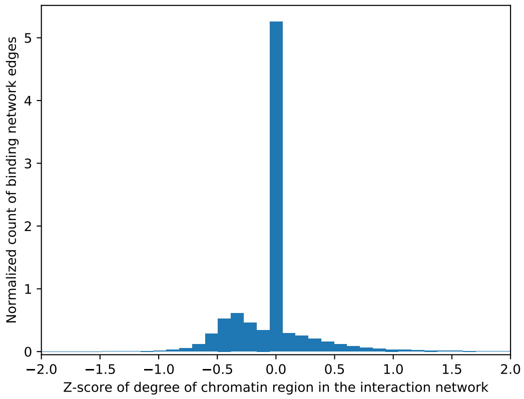


Figure S1: The distribution of z-scores of degree of regions with chip-seq peaks during our randomization process. For example, for each chip-seq peak the deviation of degree of region in the interaction network to which it is originally bound and degree of region to which it is assigned throughout 1000 randomization steps is calculated. The randomization process fairly conserves the degree of nodes.

#### Comparison with Ma et al. study

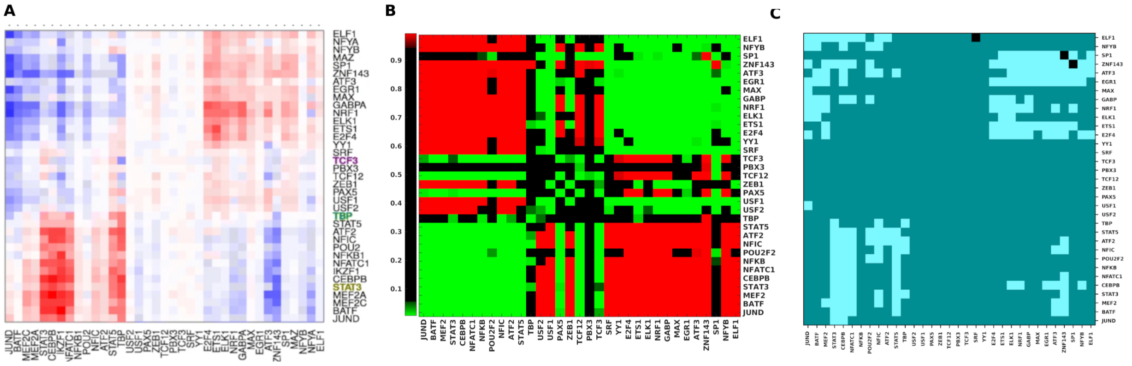


Figure S2: (A)The heatmap from the earlier study by Ma et al.[15] ((reproduced under licence CC-BY-4.0)) showing attracting TF pairs in red and repelling pairs in blue. (B) The heatmap shows the attracting and repelling pairs in green and red respectively for common TFs used in both studies, using the method we proposed in methods section. (C) The heatmap shows a qualitative comparison of the two studies as follows: bright blue = both significant, in agreement; black = both significant, in disagreement (one showing attraction, the other repulsion); dark blue = one or both insignificant.

#### Co-occurrence of various histone marks

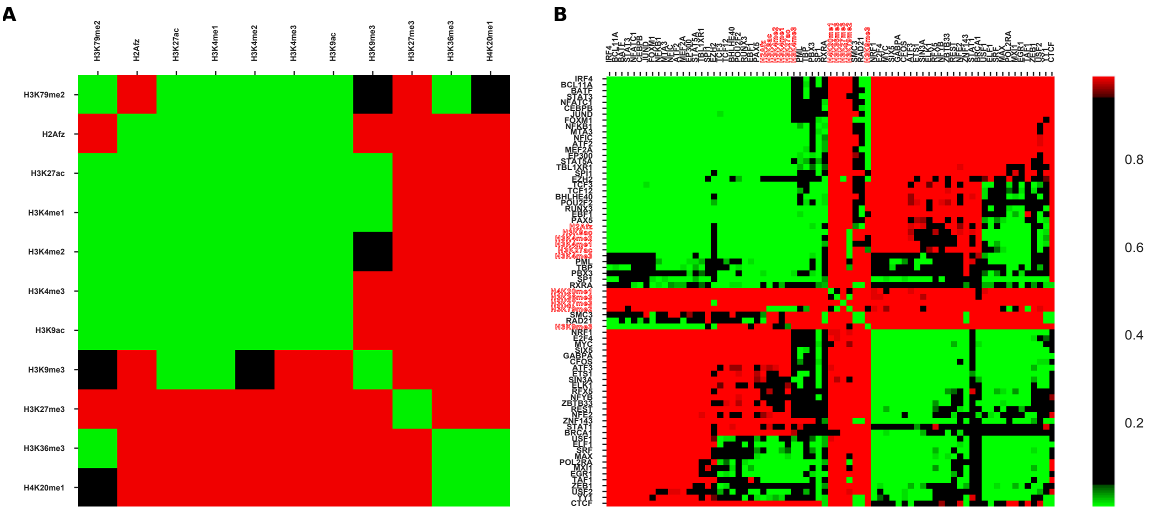


Figure S3: (A) Clustered q-value heatmap similar to figure 2 for various histone marks. (B) shows the co-occurrence pattern for all the TFs along with the histone marks (labelled in red).

#### Co-occurrence pattern in A & B compartments of the genome

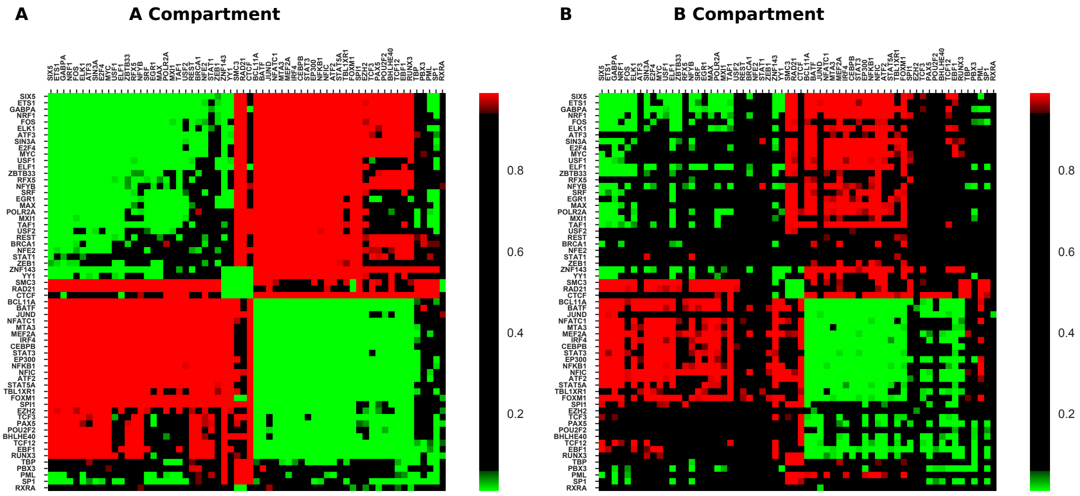


Figure S4: (A) Shows q-value heatmap of co-occurrence of TF pairs in compartment A interactions of the genome and similarly (B) shows for compartment B interactions of genome of GM12878 cell line.

#### Co-occurrence of TFBS in HeLa-S3 cell line

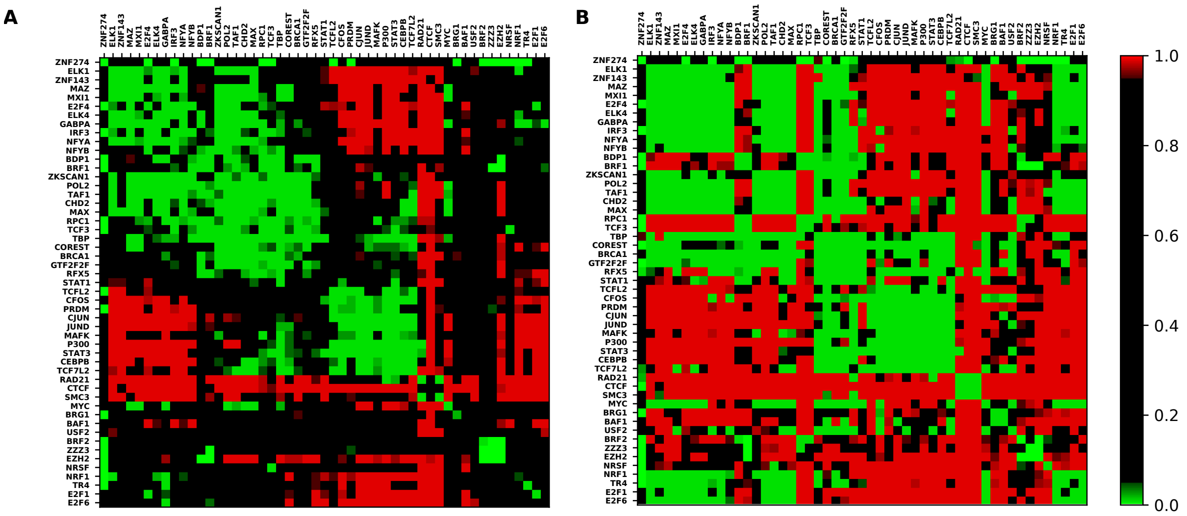


Figure S5: (A) Shows the co-occurrence of TF pairs in spatial proximal regions. (B) shows the pattern in sequential contiguous regions

#### Comparison of pattern between cell lines

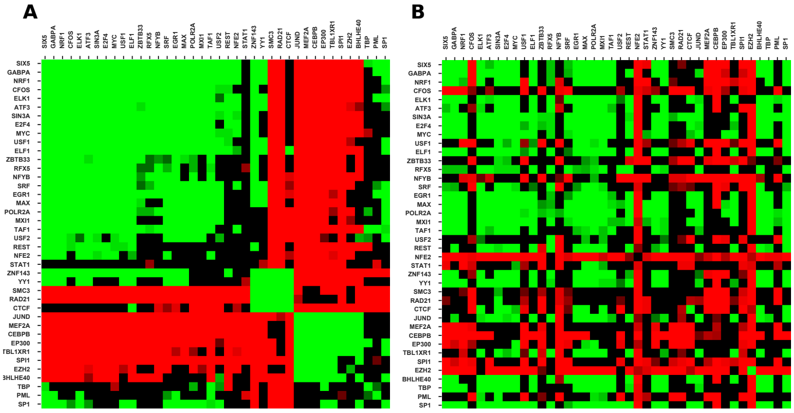


Figure S6: A comparison of co-occurrence pattern in spatial proximal regions between GM12878 and K562 cell lines for the common factors present in both. (A) shows heatmap for GM12878 cell line, (B) shows heatmap for K562 cell line.

#### Co-occurrence pattern at motif level

Motif information is available many more transcription factors than the number of factors for which ChIP-seq data is available. But several of these TFs share similar motifs. We identified clusters of similar motifs using TOMTOM[21] and selected one TF representing the clusters, which was further used in the co-occurrence analysis. The Identified clusters motifs and their logos are shown in the file `https://figshare.com/s/5285cd308c2259ac5465`.

The following table gives motifs selected as an informative motif from each cluster.

| Transcription factor | JASPAR Motif ID |
|---|---|
| Tbxt | MA0009.1 |
| ELK1 | MA0028.2 |
| Gata1 | MA0035.3 |
| Gfi1 | MA0038.1 |
| FOXI1 | MA0042.1 |
| MAX | MA0058.3 |
| NFYA | MA0060.3 |
| RXRA::VDR | MA0074.1 |
| ELK4 | MA0076.2 |
| Sox17 | MA0078.1 |
| SP1 | MA0079.3 |
| SPI1 | MA0080.4 |
| SRF | MA0083.3 |
| ZNF143 | MA0088.2 |
| TEAD1 | MA0090.1 |
| ZEB1 | MA0103.2 |
| NFKB1 | MA0105.4 |
| TP53 | MA0106.1 |
| TBP | MA0108.1 |
| ESR1 | MA0112.3 |
| NR3C1 | MA0113.3 |
| NFIC::TLX1 | MA0119.1 |
| Nkx3-1 | MA0124.2 |
| HINFP | MA0131.2 |
| STAT1 | MA0137.2 |
| REST | MA0138.1 |
| CTCF | MA0139.1 |
| Sox2 | MA0143.3 |
| Tcfcp2l1 | MA0145.1 |

| | |
|---|---|
| Myc | MA0147.1 |
| FOXA1 | MA0148.3 |
| NFATC2 | MA0152.1 |
| EBF1 | MA0154.3 |
| FOXO3 | MA0157.2 |
| EGR1 | MA0162.3 |
| CDX2 | MA0465.1 |
| DUX4 | MA0468.1 |
| E2F4 | MA0470.1 |
| ELF1 | MA0473.1 |
| Gfi1b | MA0483.1 |
| HSF1 | MA0486.2 |
| JUND | MA0491.1 |
| MAFK | MA0496.2 |
| POU2F2 | MA0507.1 |
| PRDM1 | MA0508.2 |
| RFX5 | MA0510.2 |
| RUNX2 | MA0511.2 |
| Rxra | MA0512.2 |
| Tcf12 | MA0521.1 |
| Esrra | MA0592.2 |
| Hoxa9 | MA0594.1 |
| SREBF1 | MA0595.1 |
| FOXG1 | MA0613.1 |
| Mitf | MA0620.1 |
| mix-a | MA0621.1 |
| BARHL2 | MA0635.1 |
| BHLHE41 | MA0636.1 |
| ETV6 | MA0645.1 |
| GRHL1 | MA0647.1 |
| IRF9 | MA0653.1 |
| NFIA | MA0670.1 |
| NKX2-3 | MA0672.1 |
| ONECUT1 | MA0679.1 |
| PAX7 | MA0680.1 |
| POU4F2 | MA0683.1 |
| SP4 | MA0685.1 |
| ZBTB7B | MA0694.1 |
| OTX2 | MA0712.1 |
| GLIS1 | MA0735.1 |
| GLIS3 | MA0737.1 |
| Hic1 | MA0739.1 |
| SCRT2 | MA0744.1 |
| E2F7 | MA0758.1 |
| Tcf7 | MA0769.1 |
| MEF2D | MA0773.1 |
| MEIS3 | MA0775.1 |
| PAX1 | MA0779.1 |
| PAX9 | MA0781.1 |
| POU3F4 | MA0789.1 |
| SMAD3 | MA0795.1 |
| TGIF1 | MA0796.1 |
| TFAP2C(var.3) | MA0815.1 |
| ATF7 | MA0834.1 |
| NFE2 | MA0841.1 |
| TEF | MA0843.1 |
| Rarb | MA0857.1 |
| Rarg(var.2) | MA0860.1 |
| E2F2 | MA0864.1 |

| | |
|---|---|
| SOX4 | MA0867.1 |
| Sox11 | MA0869.1 |
| TFAP2A(var.3) | MA0872.1 |
| HOXC13 | MA0907.1 |
| SIX1 | MA1118.1 |

Table S3: The selected motifs after clustering for similar motifs using TOMTOM

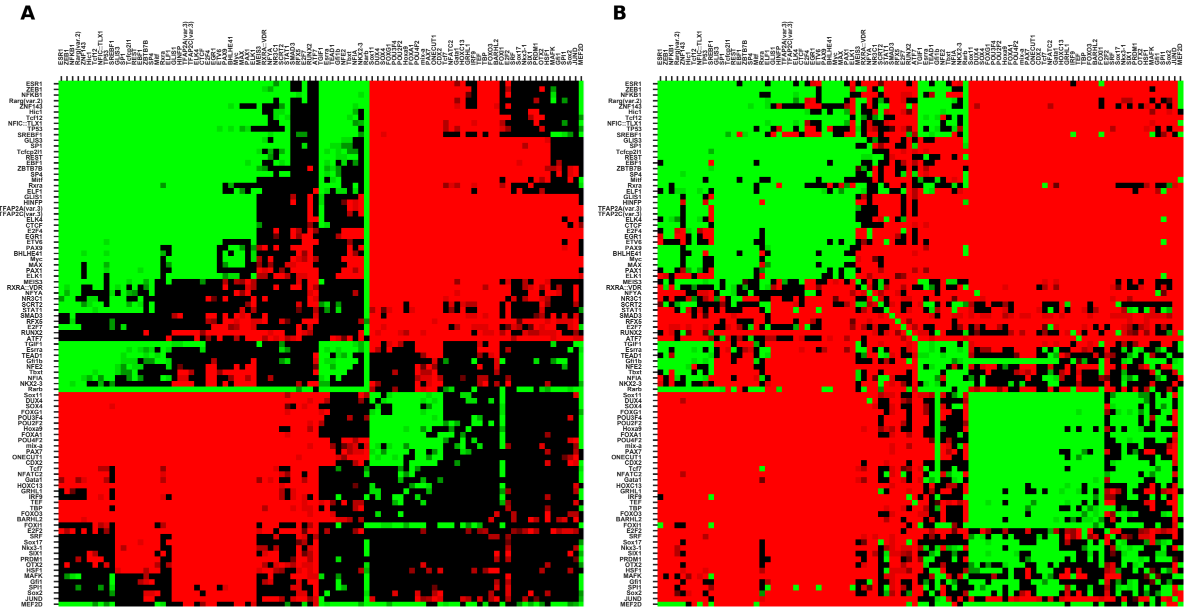


Figure S7: (A) The co-occurrence of TF motifs sites in spatial proximal regions and (B) in sequential contiguous regions of GM12878 cell line

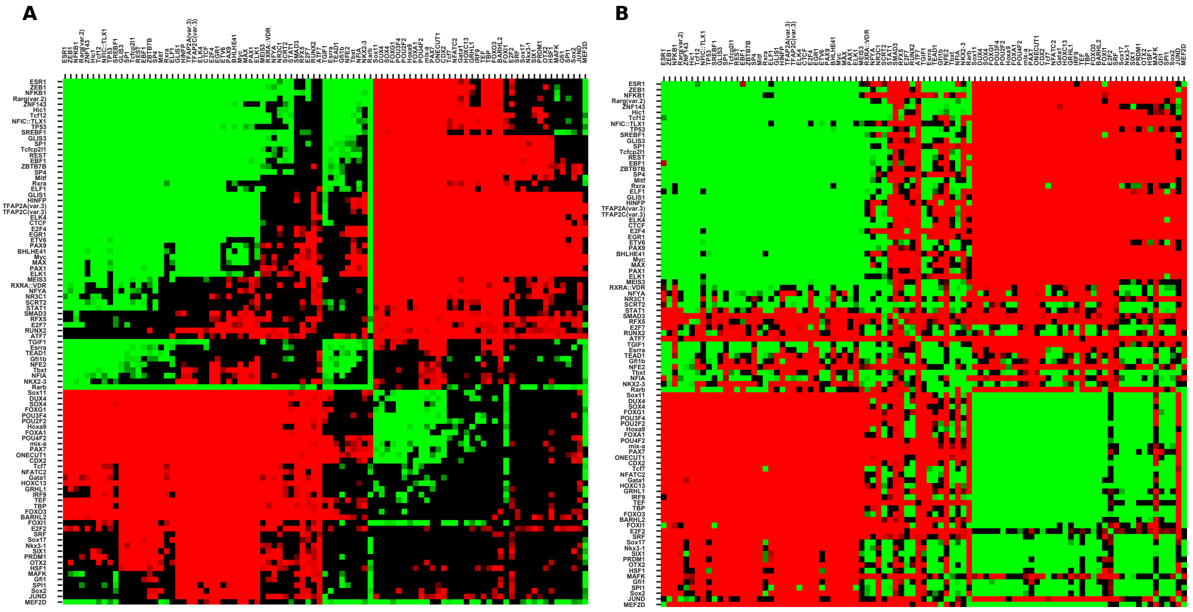


Figure S8: (A) The co-occurrence of TF motifs sites in spatial proximal regions and (B) in sequential contiguous regions of k562 cell line

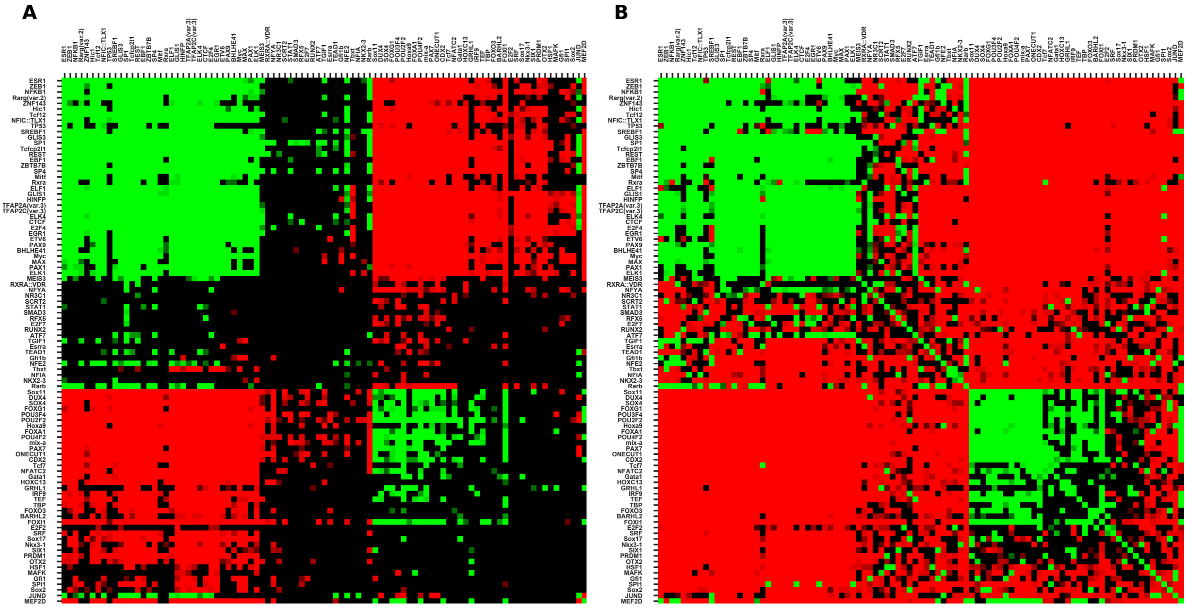


Figure S9: (A) The co-occurrence of TF motifs sites in spatial proximal regions and (B) in sequential contiguous regions of HeLa-S3 cell line

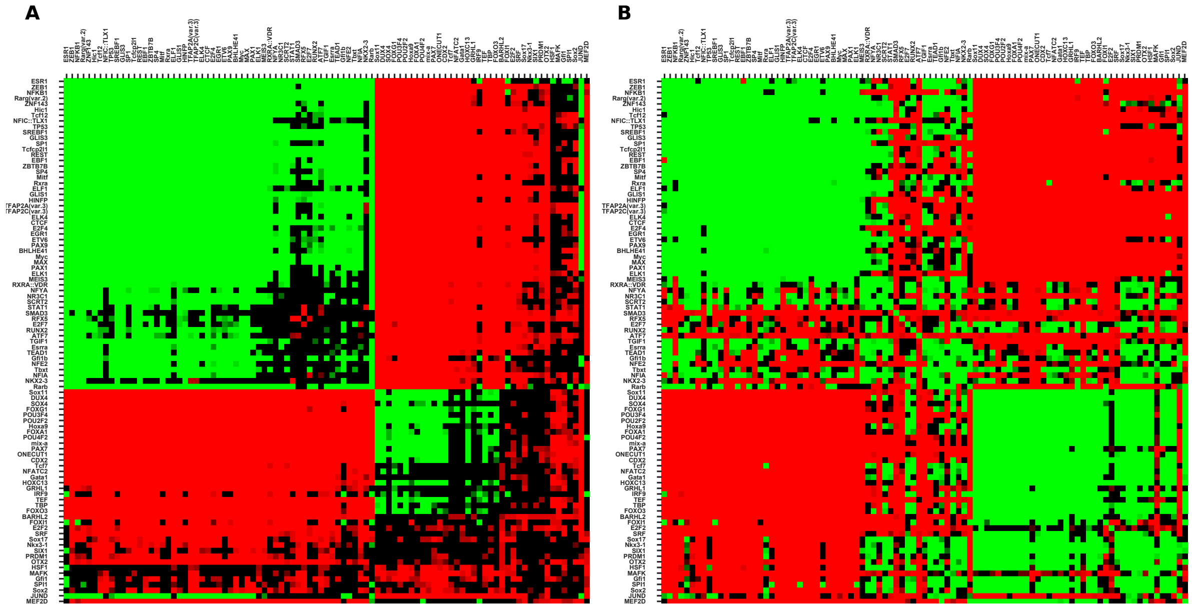


Figure S10: (A) The co-occurrence of TF motifs sites in spatial proximal regions and (B) in sequential contiguous regions of MCF7 cell line

### Comparison of all four types of co-occurrence patterns

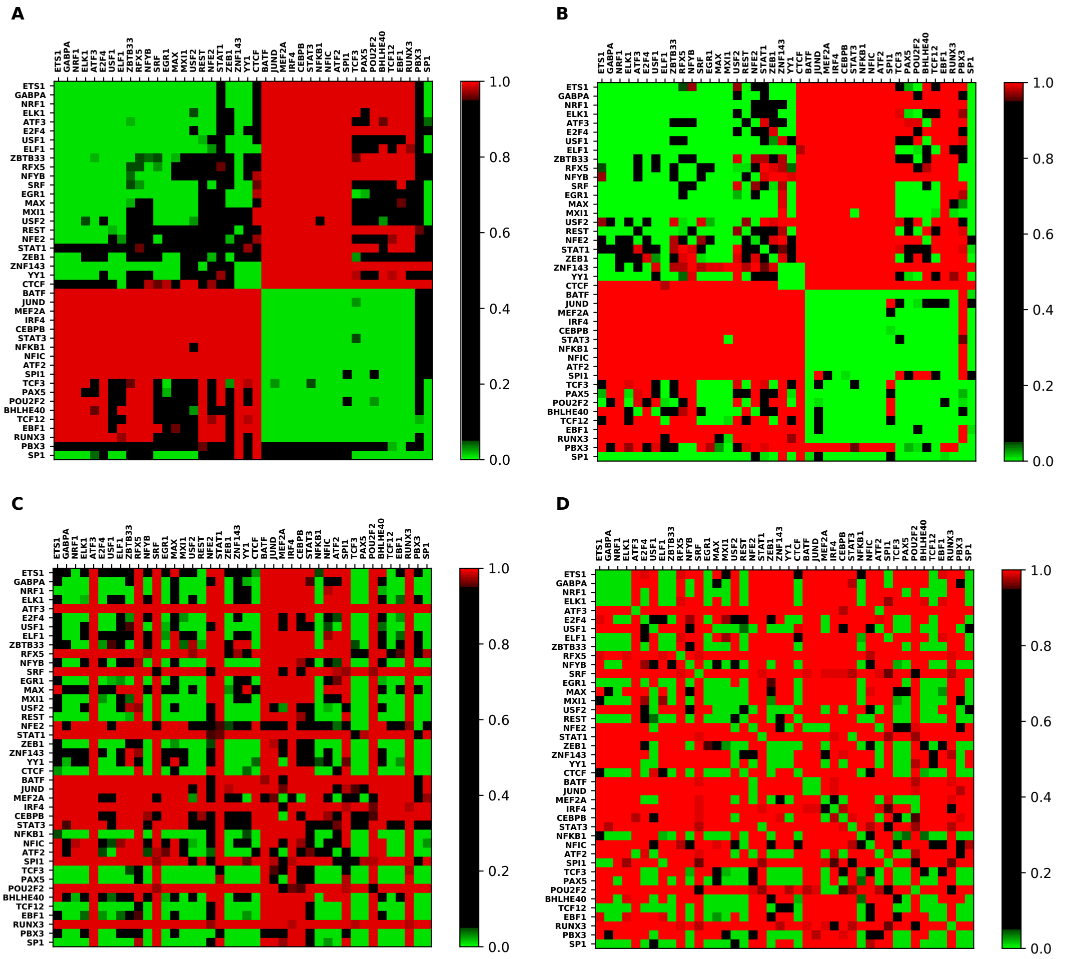


Figure S11: The figure shows all the four different q-value heatmaps of GM12878 cell line. (A) using TF chip-seq peaks in spatial proximal regions, (B) using TF chip-seq peaks in sequential contiguous regions, (C) using TF motif sites in spatial proximal regions, and (D) using TF motif sites in sequential contiguous regions.

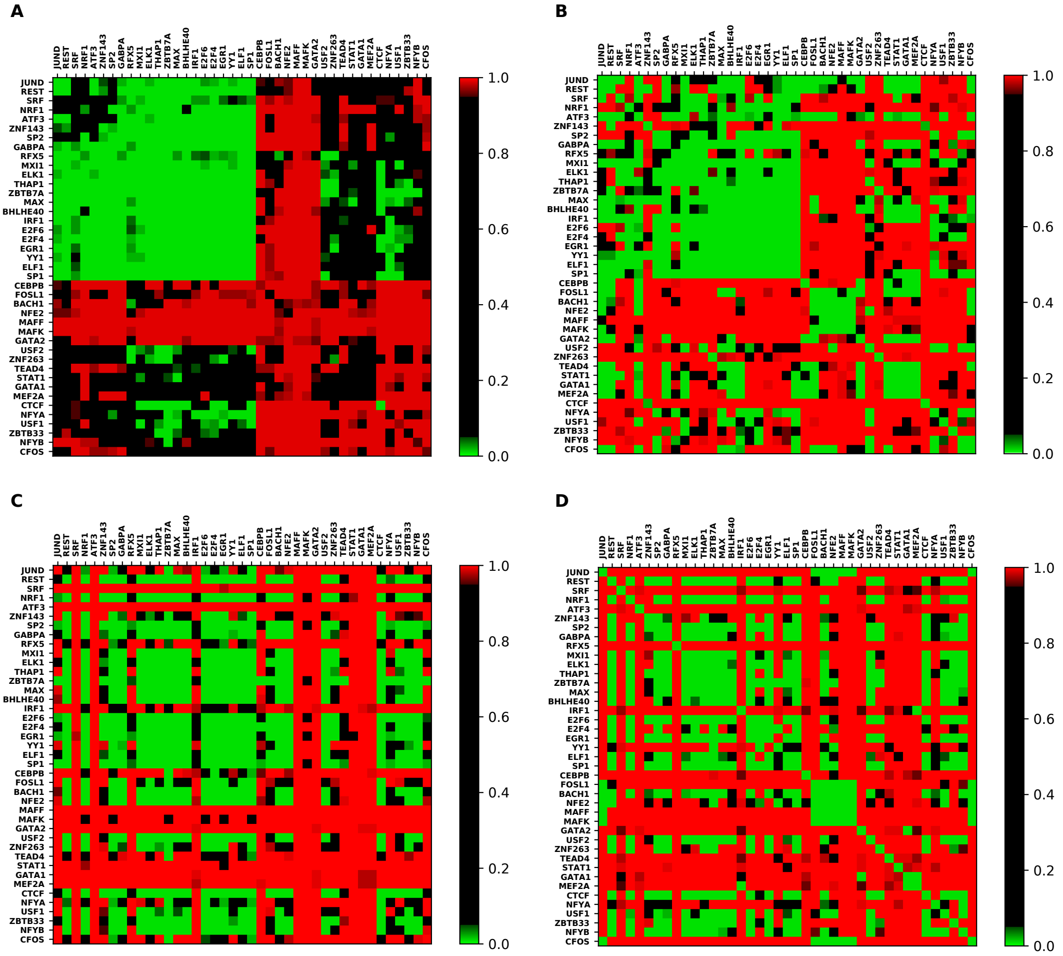


Figure S12: The figure shows all the four different q-value heatmaps of K562 cell line. (A) using TF chip-seq peaks in spatial proximal regions, (B) using TF chip-seq peaks in sequential contiguous regions, (C) using TF motif sites in spatial proximal regions, and (D) using TF motif sites in sequential contiguous regions.

#### Motif strength analysis

| Transcription factor | JASPAR motif ID |
|---|---|
| ETS1 | MA0098.3 |
| GABPA | MA0062.1 |
| NRF1 | MA0506.1 |
| ELK1 | MA0028.2 |
| ATF3 | MA0605.2 |
| E2F4 | MA0470.2 |
| USF1 | MA0093.2 |
| ELF1 | MA0473.2 |
| ZBTB33 | MA0527.1 |
| RFX5 | MA0510.2 |
| NFYB | MA0502.2 |
| SRF | MA0083.1 |
| EGR1 | MA0162.3 |
| MAX | MA0058.1 |
| MXI1 | MA1108.1 |
| USF2 | MA0526.1 |
| REST | MA0138.2 |
| NFE2 | MA0841.1 |
| STAT1 | MA0137.3 |

| | |
|---|---|
| ZEB1 | MA0103.2 |
| ZNF143 | MA0088.2 |
| YY1 | MA0095.2 |
| CTCF | MA0139.1 |
| BATF | MA1634.1 |
| JUND | MA0491.1 |
| MEF2A | MA0052.3 |
| IRF4 | MA1419.1 |
| CEBPB | MA0466.2 |
| STAT3 | MA0144.2 |
| NFKB1 | MA0105.2 |
| NFIC | MA0161.2 |
| ATF2 | MA1632.1 |
| SPI1 | MA0080.4 |
| TCF3 | MA0522.2 |
| PAX5 | MA0014.2 |
| POU2F2 | MA0507.1 |
| BHLHE40 | MA0464.2 |
| TCF12 | MA1648.1 |
| EBF1 | MA0154.3 |
| RUNX3 | MA0684.1 |
| PBX3 | MA1114.1 |
| SP1 | MA0079.4 |

Table S4: This table gives the JASPAR motif ids for the TF used in the motif strength analysis in Figure 9

#### Target gene analysis

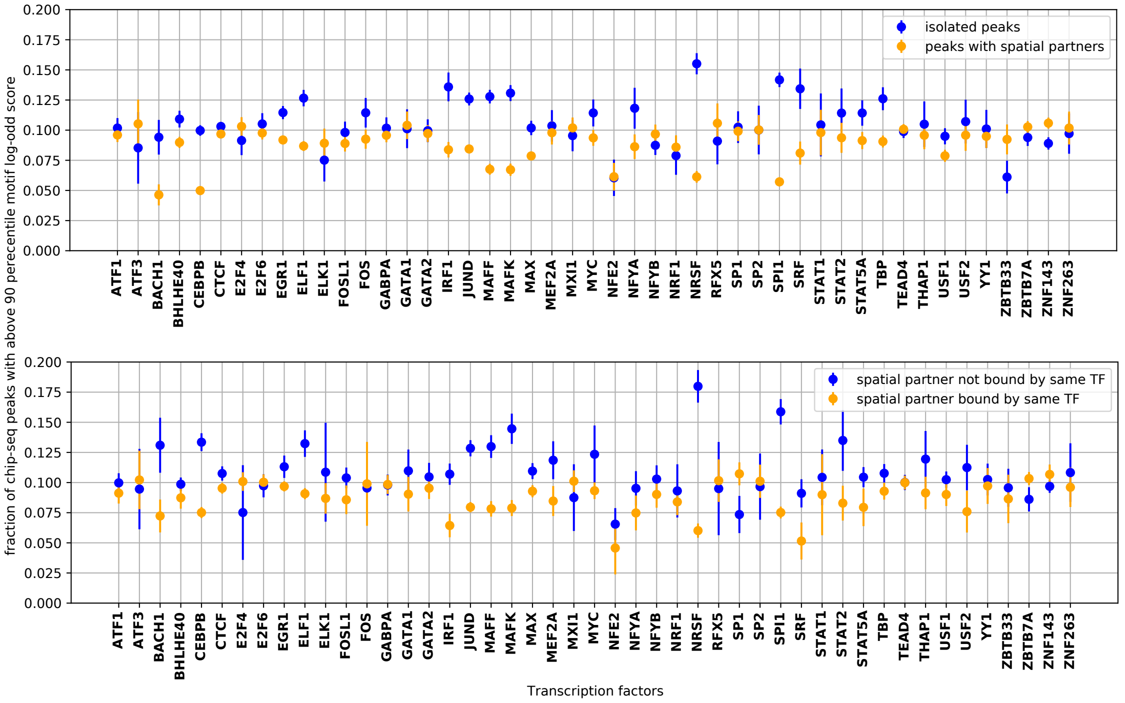


Figure S13: (A) Spatially isolated ChIP-seq peaks have a higher fraction of strong motifs than peaks with spatial clusters . (B) Even among the peaks with spatial partners, the peaks with the partner region not bound by same TF show higher fraction of strong motifs than bound by the same TF.

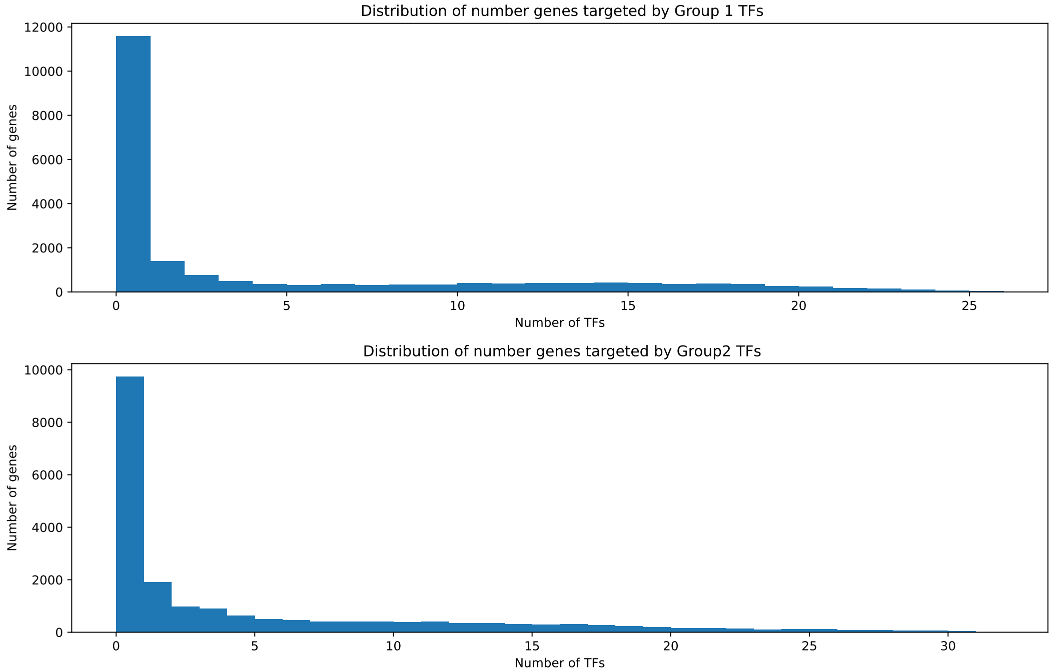


Figure S14: Distribution of number of putative TF regulators per gene in GM12878 cell line.

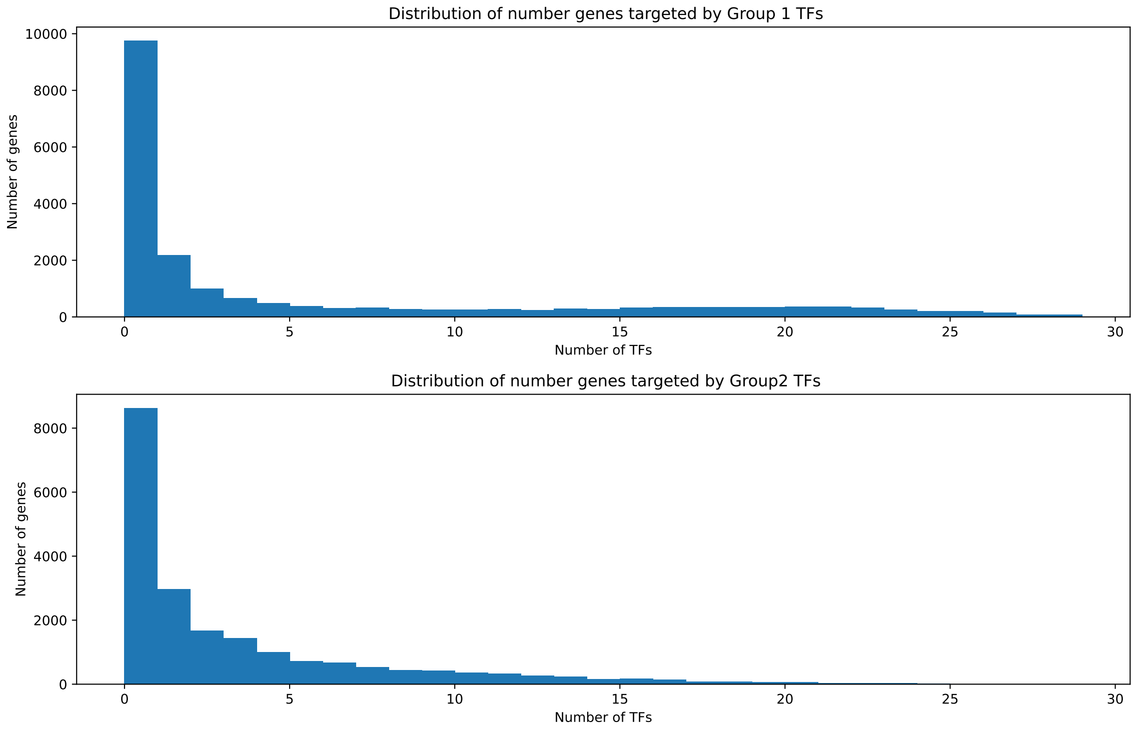


Figure S15: Distribution of number of putative TF regulators per gene in GM12878 cell line.

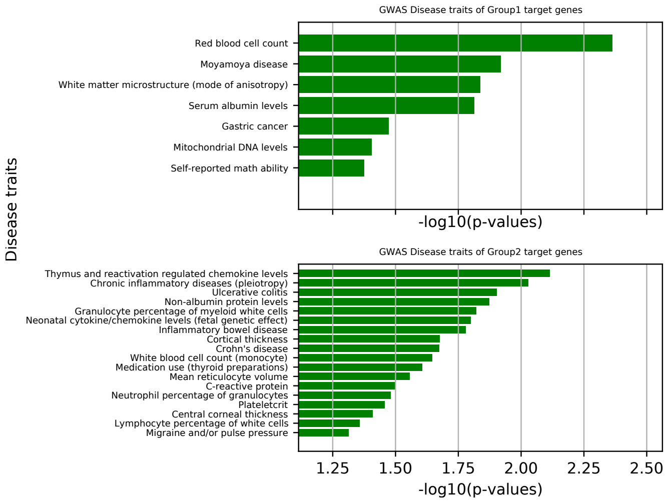


Figure S16: The enriched GWAS disease traits of the target genes of Group 1 and Group 2 TFs

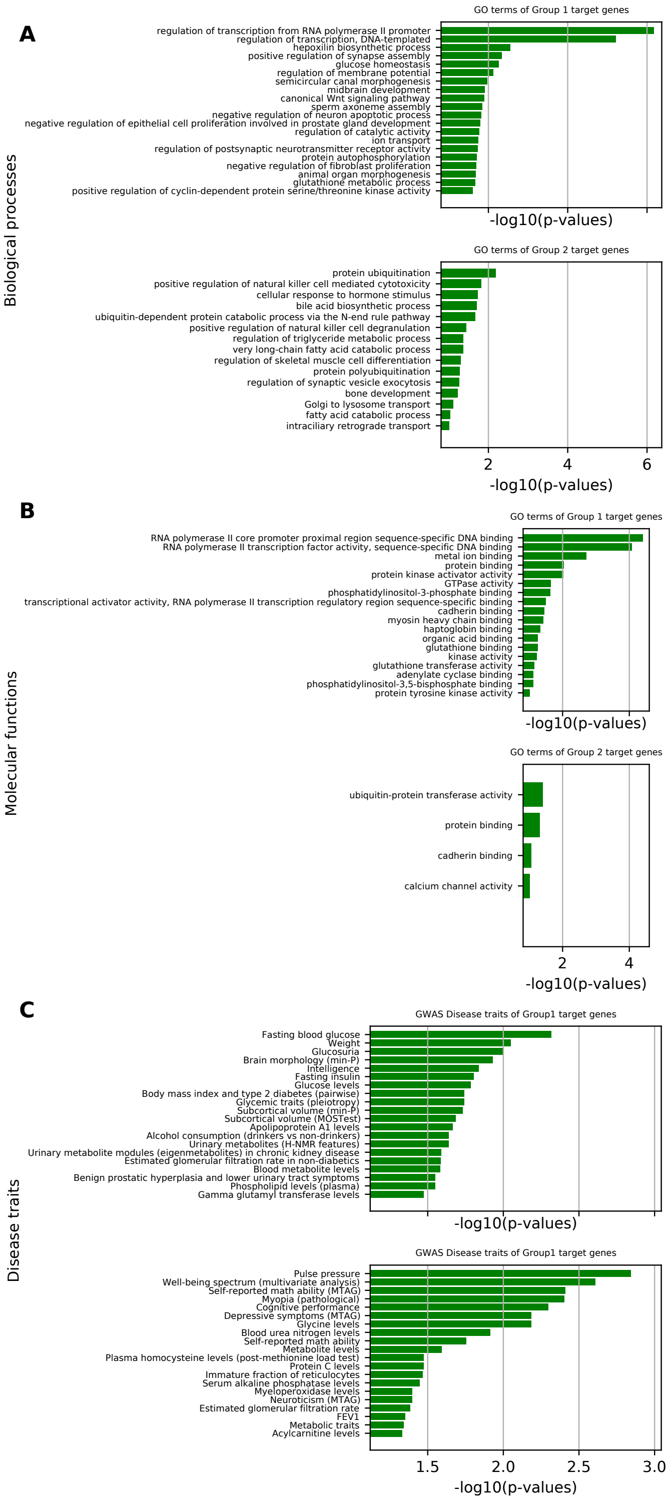


Figure S17: The enriched GO biological processes, molecular function, and GWAS disease trait terms for the Group 1 and the remaining TFs used in the study for the K562 cell line

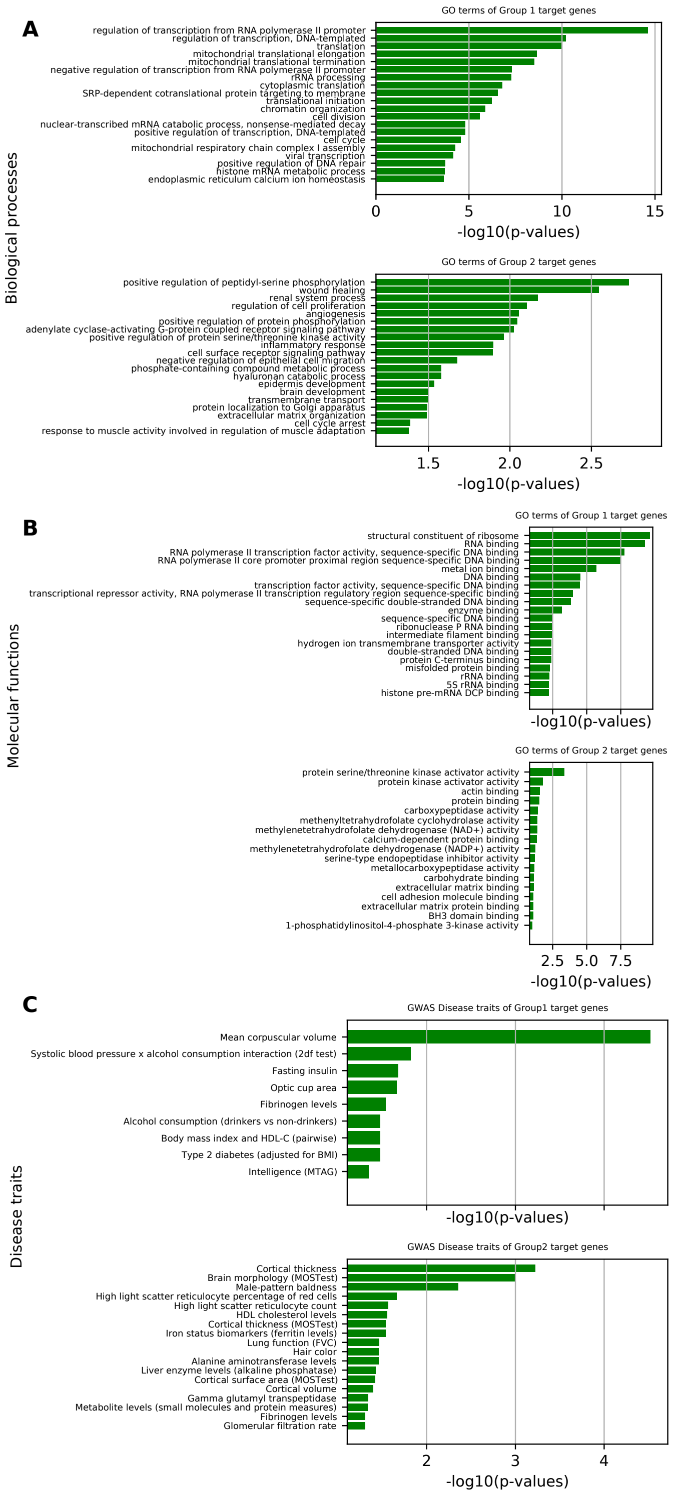


Figure S18: The enriched GO biological processes, molecular function, and GWAS disease trait terms for the Group 1 and Group 2 TFs used in the study for the HeLa-S3 cell line
